# High-resolution genome and genetic map of tetraploid *Allium porrum* expose pericentromeric recombination

**DOI:** 10.1101/2025.04.21.649809

**Authors:** Ronald Nieuwenhuis, Roeland Voorrips, Danny Esselink, Thamara Hesselink, Elio Schijlen, Paul Arens, Jan Cordewener, Olga Scholten, Sander Peters

## Abstract

We present the first reference genome of highly heterozygous autotetraploid *Allium porrum* (leek). Combining long-read sequencing with SNP-array screening of two experimental F1 populations, we generated a genetic map with 11,429 SNP markers across 8 linkage groups and a chromosome-scale assembly of *Allium porrum* (leek) totaling 15.2 Gbp in size. High quality of the reference genome is substantiated by 97.2% BUSCO completeness and a mapping rate of 96% for full-length transcripts. The linkage map exposes the recombination landscape of leek and confirms that crossovers are predominantly proximal located to the centromeres, contrasting with distal recombination landscapes observed in other *Allium* species. Comparative genomics revealing structural rearrangements between *A. porrum* and its relatives (*A. fistulosum*, *A. sativum*, *A. cepa*), suggests a closer genomic relationship to *A. sativum*. Our annotated high-quality reference genome delivers crucial insights into the leek genome structure, recombination landscape, and evolutionary relationships within the *Allium* genus, offering significant implications for breeding programs, facilitating marker-assisted selection and genetic improvement in leek.

## Introduction

Leek (*Allium ampeloprasum* var. *porrum* syn. *Allium porrum*) is a member of the *Allium* genus to which also the well-known onion (*A. cepa*), shallot (*A. cepa* var. *aggregatum*), garlic (*A. sativum*), Japanese bunching onion or Welsh onion (*A. fistulosum*), scallion (*A. cepa* var. *cepa*), chives (*A. schoenoprasum*) and Chinese onion (*A. chinense*) vegetables belong. Historically, Allium species have been cultivated and appreciated for consumption already from the second millennium BC onward, as evidenced by dried specimens found from archeological sites in Egypt and Mesopotamia (Hehn, 1870; Zohary *et al*., 2012) and documented Roman recipes (Sanderson & Renfrew, 2005). Today, cultivated leek is globally appreciated for its mild onion-like taste and nutritional value with high content of vitamin B6, C and K, folate, iron and manganese.

Leek is a cross-fertilizing species and is generally considered a tetraploid (2n=4x=32) (Kik *et al*., 2021). Both auto- and (weak) segmental allopolyploidy are reported in different studies based on varying levels of multivalent formation (Levan, 1940; Kadry & Kamel, 1955; Murín, 1964; Koul & Gohil, 1970). Although leek can self-fertilize to a limited extent of 20%, it displays strong inbreeding depression with plant-weight reductions of up to 60% (Berninger & Buret, 1967). In general, modern leek breeding concentrates on better crop uniformity, higher yield and pest and disease resistance. To obtain better uniformity and higher yield, mostly F1-hydrids are exploited as commercial varieties. Because of intensive selection by leek growers since the last century, a significant amount of genetic variation originally present in the old landraces has been lost in leek germplasm. The genetic erosion has further challenged leek breeders for the identification and introgression of resistance to leek moth *(Acrolepiopsis assectella),* leek fly *(Delia antiqua),* thrips *(Thrips tabaci*), onion leaf miner (*Liriomyza cepae)* stem and bulb nematodes (*Ditylenchus dipsaci, Meloidogyne* spp., *Pratylenchus penetrans, Paratrichodorus* spp., *Trichodorus* spp. and *Longidorus* spp.), smudge (*Colletotrichum circinans),* leaf blotch *(Cladosporium allii),* white tip disease *(Phytophthora porri),* purple blotch *(Alternaria porri),* leek rust *(Puccinia allii),* basal rot *(Fusarium culmorum),* white rot *(Sclerotium cepivorum),* black mould *(Pleospora herbarum),* bacterial diseases caused by *Pseudomonas* spp. and *Erwinia spp.,* and aphid transmitted viral diseases such as Leek Yellow Stripe Virus (Becue, 2003; Lorbeer *et al*., 2002; Mark *et al*., 2002; Salomon, 2002). Until today resistance to many of these diseases and pests in cultivated leek crops is absent.

For genetic improvement of Allium crops, crop wild relatives (CWR) are considered important genetic resources (Kik *et al*., 2021). Despite successful use of introgression breeding with cross-fertilizing wild species to transfer beneficial alleles in many target crops, Allium breeders have faced significant challenges in using wide hybridization for leek crop improvement. The success of introgression breeding depends on the biological process known as meiotic recombination, a biological process by which genetic material from the donor into the recipient crop is introduced. The exchange of genetic material between maternal and paternal chromosomes is commonly referred to as crossover (CO). As observed for many organisms, also in Allium species COs are not randomly distributed, and their distribution and localization appear genetically and strictly controlled (Kudryavtseva *et al*., 2023). Furthermore, the designation of COs varies considerably between Allium species. Recently, Kudryavtseva and coworkers (2023) demonstrated that *A. fistulosum* COs predominantly occur in the proximal chromosome regions, while in *A. cepa,* COs mainly manifest in the distal and interstitial chromosome regions. These differences may result in homoeologous chromosome pairing disorders during meiosis in hybrids, leading to reduced COs in regions containing beneficial alleles thus hampering introgression breeding. Indeed, Allium introgression lines often display genome instability through a phenomenon known as genome dominance, manifested by unequal inheritance and loss of introgressed genetic donor material in successive generations, further impeding the employment of wide hybridization in practical Allium breeding (Kopecký *et al*., 2022). Thus, understanding of the recombination and CO landscapes is crucial for successful introgression breeding in Allium crops.

The development of a reference genome and genetic map offers a significant opportunity to overcome many of these challenges in leek breeding. These tools enable the assessment of genetic variation and inheritance, facilitating more efficient breeding strategies. A major advantage is the improved ability to identify beneficial haplotypes for disease resistance. Given leek’s widespread susceptibility to pests, fungi, bacteria, and viruses, distinguishing between parental haplotypes in an outcrossing autotetraploid would allow breeders to track, select, and monitor the retention of specific resistance alleles over generations. Additionally, linkage maps help clarify recombination patterns and CO distribution. Since COs are not randomly distributed, a genome assembly and genetic map provide insight into CO patterns, allowing breeders to select compatible breeding lines for successful introgression breeding. This is particularly helpful when incorporating alleles from CWR and avoiding homoeologous chromosome pairing issues (Jones *et al*., 1996; Khazanehdari & Jones, 1997; Kiełkowska, 2012). Linkage maps also help identify and track deleterious recessive alleles that may negatively affect plant fitness, aiding breeders in selecting complementary parental lines with optimized agronomic performance and disease resistance, while minimizing inbreeding depression. Furthermore, structural variation detection is another critical advantage of reference genomes and genetic maps. Large-scale genomic variations, such as copy number variations, inversions, and translocations, can influence key agronomic traits. By combining detailed genome sequences with genetic map data, breeders can better understand these variations and selectively retain or eliminate structural variants that impact crop performance. Genetic maps also play a crucial role in stabilizing introgression lines, allowing the effective transfer of beneficial traits by tracking wild allele inheritance and identifying regions prone to instability.

The sheer size, autotetraploid heterozygous nature, and highly repetitive structure of the leek genome have long posed challenges for genome assembly and genetic map construction. In this paper, we present the construction of the first chromosome-scale reference genome for leek, along with the development of a genetic map, marking a significant milestone in Allium genomics and plant breeding. By resolving the complexities of this genome at a chromosome scale, this study provides a foundation for dissecting genetic variation, understanding recombination landscapes, and accelerating breeding efforts. The ability to phase haplotypes and map crossovers will allow for more precise tracking of beneficial alleles, especially in the context of introgression breeding, where homoeologous recombination, chromosome synapsis issues, and genome instability have previously hindered progress (Jones *et al*., 1996; Khazanehdari & Jones, 1997; Kiełkowska, 2012; Kopecký *et al*., 2022). Moreover, the high level of genomic resolution enables the identification of structural variants that influence agronomic traits, facilitating targeted selection strategies. The reference genome and genetic map presented here are vital resources for leek improvement, set a precedent for genomic studies in complex Allium species, and complement ongoing efforts in onion, garlic, and Welsh onion (Sun *et al*., 2020; Hao *et al*., 2023). Ultimately, this achievement paves the way for more resilient, genetically diverse, and high-performing leek cultivars, ensuring the sustainable advancement of this globally important crop.

## Materials and methods

### Plant material

We established a genome sequence from an individual leek plant of accession “Leidse prei 2018-94”, a landrace/grower selection from the Netherlands. Linkage mapping was performed in two populations: one F1 of the cross 34012-7 (a genic male sterile (gms) plant donated by Bejo Zaden, the Netherlands) x the “Leidse prei 2018-94” plant mentioned above, consisting of 187 plants, and one F1 of the cross 17169-29 (a gms plant donated by Nunhems / BASF crop science, the Netherlands) x *A. ampeloprasum*, consisting of 248 plants.

### DNA isolation & library prep

Seven clones of the leek father plant “Leidse prei 2018-94" were used for tissue harvest. Multiple young most inner leaves were harvested, snap frozen in liquid nitrogen and stored at -80°C. Approximately 1.5 g of leaf material was ground with liquid nitrogen and used for HMW gDNA isolation using NucleoBond HMW DNA kit following manufacturer’s instructions (Machery-Nagel). Additionally, 1 g of leaf material was ground with liquid nitrogen and used for HMW gDNA isolation using MagMAX Plant DNA kit following manufacturer’s instructions (Applied Biosystems). DNA quantity, purity and fragment size was determined using Qubit (Invitrogen), OD values (NanoDrop) and Fragment Analyzer (Agilent). Obtained gDNA was sheared in different rounds by Megaruptor 2 (Diagenode) with 20 to 30 kbp target fragment size. Small fragments were removed by utilizing Blue Pippin (Sage Science) for size selection. Size selected DNA was used for six SMRTbell library preps using SMRTbell Express Template Preparation kit 2.0 according to manufacturer’s instructions (Pacbio; Procedure-Checklist-Preparing-HiFi-SMRTbell-Libraries-using-SMRTbell-Express-Template-Prep). SMRTbell libraries were subjected to DNA polymerase complexing using Sequel binding kit v2.0/3.0 and Sequel Polymerase 3.0/2.2. Final sequencing was done using sequence primer v4 /v5 on a Pacbio Sequel-II/ Sequel-IIe instrument (see section Genome sequencing). Sequencing reactions were performed with sequencing kit 2.0 /3.0, using adaptive loading, 65 to 85 pM on plate loading concentration, 120- or 240-minutes immobilization time and 30 hours movie time per SMRT cell.

### Genome sequencing

Long insert libraries of 15 and 19 kbp were sequenced on the PacBio Sequel-II(e) platform using 44 SMRT cells (Supplementary Table S1). Subsequent circular consensus sequences (CCS) were called using the pbccs v5.0.0 command line utility. HiFi reads were defined as CCS reads having a minimum number of 3 passes and a mean read quality score of Q20. Quality control per SMRTcell was checked via SMRTlink and an in-house pipeline including FastQC v0.11.9 (Andrews, 2010), KMC v3.1.1 (Kokot *et al*., 2017), Smudgeplot v0.2.3dev_rn (Ranallo-Benavidez *et al*., 2020), GenomeScope v2 (Ranallo-Benavidez *et al*., 2020) and BlastN v2.11.0+ (Altschul *et al*., 1990; Camacho *et al*., 2009) with the NCBI nt, plastid and mitochondrion publicly available databases downloaded 2021-04-02 (Sayers *et al*., 2024). Reads produced from different libraries were then combined into a single data set for further analyses.

### Contig assembly

Combined reads of all sequenced libraries were assembled using hifiasm v0.15.1-r334 (Cheng *et al*., 2021) with output settings for primary and alternative assembly selected. All contigs from both the primary and alternative assemblies were screened for contamination using NCBI Foreign Contamination Screen v0.4.0 (Astashyn *et al*., 2024), and non-subject sequences were subsequently removed. Splitting of contigs was done with NCBI Adaptor v0.4.0 in case of detected adaptor sequences. Any remaining adaptors sequences were removed to avoid assembly errors. BUSCO v5.2.2 (Manni *et al*., 2021a, 2021b; Simão *et al*., 2015) was used with the embryophyte odb10 database (2020-09-10) to check completeness of conserved single copy orthologs. Further purging of the primary assembly was performed in a customized manner to accommodate the large dataset and limited sequence coverage. The method optimized single copy BUSCO gene percentage in a minimal set of contigs. (Supplementary Figure S1).

Selected contigs were used for subsequent annotation and scaffolding. K-mer based completeness checks were omitted due to the heterozygous and autopolyploid nature of the sequenced specimen, with the purged result representing only a pseudo-haploid version of the genome.

Reference based assembly was applied to the leek WGS HiFi dataset to assemble the mitochondrion and chloroplast genomes. For the plastid genome ptGAUL v1.0.5 (Zhou *et al*., 2023) was used with the A. *ampeloprasum* reference sequence (NC_044666.1). For the mitochondrial genome mitoHIFI v3.0.1 (Uliano-Silva *et al*., 2023) and ptGAUL v1.0.5 assemblers were used with reference from *A. cepa* and *A. sativa*.

### RNA isolation & library prep

For RNA isolation, material of different tissues (green leaf, stem section green / white, basal plate section, roots) from leek father plant (2018-94-02) were collected. In addition, flower tissue was collected from a different plant (leek mother plant 1716929 ms), immediately snap frozen in liquid nitrogen, and stored at -80°C. RNA from the father plant tissues was isolated using Ambion PureLinkRNA Mini Kit (Life Technologies). RNA from the mother plant flower tissue was isolated using ZymoBIOMICS RNA Mini prep Kit (Zymo Research). Subsequently, RNA quantity and purity was analyzed by Qubit (Invitrogen), OD values (NanoDrop) and Bioanalyzer RNA plant pico assay (Agilent).

Of each tissue, 300 ng total RNA was used to create a barcoded IsoSeq library following manufacturer’s guidelines (Pacbio; Procedure-Checklist-Iso-Seq-Express-Template-Preparation-for-Sequel-and-Sequel-II-Systems). SMRTbell library yield was quantified by Qubit (Invitrogen) and SMRTbell sizes were checked by Bioanalyzer High sensitivity DNA Assay (Agilent). Libraries were pooled equimolar, subjected to DNA Polymerase SMRTbell complexing using Sequel II Binding kit 2.0. and primer v4 prior to loading on 4 SMRT cells with 50 to 58 pM on plate loading concentration. Sequencing reaction was performed on a PacBio Sequel-II system with 24-hour movie time.

### Transcriptome sequencing

Genetic diversity was mined for array probe design by using RNA-seq of 14 *A. porrum* accessions including the mapping population parents “Leidse prei 2018-94”, MS34012-7, and species *A. lusitanicum* and *A. eduardii* (Supplementary Table S2). RNA-seq libraries of young plantlets and leaf tissue were sequenced on an Illumina Novaseq6000 platform using an S2 flow cell. Base calling and initial quality filtering of raw sequencing data was done with bcl2fastq v2.20.0.422 (https://emea.support.illumina.com/downloads/bcl2fastq-conversion-software-v2-20.html) using default settings.

Next, full-length transcripts from different tissues of the sequenced subject were sequenced, using PacBio IsoSeq on a Sequel-IIe platform for annotation of the *de novo* assembled genome. Consensus reads were called with the ccs v5.0.0 command-line utility of PacBio. HiFi reads were generated using the same specifications as used for the genomic reads. Reads were then stripped of primers and demultiplexed using lima v2.0.0 (https://github.com/PacificBiosciences/barcoding). Poly-A tails were trimmed and concatemers were removed using isoseq3 v3.4.0 (https://github.com/PacificBiosciences/IsoSeq) to generate full length non-concatemer reads, which were subsequently clustered using isoseq3 cluster (https://github.com/ylipacbio/IsoSeq3) without final polishing.

### Axiom array design

To create a genetic map, first a reference transcriptome sequence set was constructed by clustering all the IsoSeq data of “Leidse prei 2018-94” with isONclust (Sahlin & Medvedev, 2020). This reference formed the basis for variant calling. A set of largest 45K clusters was then selected based on the completeness and duplication rate of the BUSCO score (74% completeness with 10% duplication rate). The final reference sequence set, consisting of the largest 45K isONclust sequences, was concatenated with the chloroplast genome of *A. ampeloprasum* to accommodate subsequent read mapping and filtering (Filyushin *et al*., 2019). IsoSeq reads of samples 2018-94-02 and MS17169-29-2 were mapped using Minimap2 v2.11-r797 (Li, 2018) with additional settings (splice, -secondary=no, -C5 -06,24, -B4). Illumina reads of samples MS 17169-29-2, MS 34012-7 and 2018-94-1-1 were mapped using STARv020201 (Dobin *et al*., 2013) with default settings. Duplicate read pairs were marked with Picard tools (v2.2.1) MarkDuplicates (https://broadinstitute.github.io/picard/). The bam files were screened with samtools (Danecek *et al*., 2021) to filter for non-primary and supplementary alignments (sam flag -F2304). NGSEP3 v4.01 (Tello *et al*., 2019) MultisampleVariantsDetector was used for variant calling of the parents of the two segregating populations (default settings with -ploidy=4). Biallelic SNP variants were selected having a flanking sequence of at least 30 bases without additional SNPs or indels. Probes with a low complexity sequence, low GC-content, having an A/T or C/G variant, or showing redundancy, were subsequently discarded. For the design of the SNP array, three priorities were defined: Priority 1 focused on probes that needed to be called in both sets, Priority 2 prioritized probes called in the parents of the first cross, and Priority 3 prioritized probes in the parent of the second cross. Following the probe design, a draft *de-novo* assembly of a non-purged genome became available that was used to filter the recommended probes. For that the mapped probes were screened with Bowtie2 (--very sensitive) (Langmead & Salzberg, 2012) over possible intron/exon boundaries. Probes with more than the expected maximum of 4 possible hits in a tetraploid individual were filtered out. maximum number of 350 probes per contig were chosen.

### Dosage calling

The array hybridization was performed in two separate experiments, one with each of the two F1 populations, along with replicate samples of the parents and in the case of the second F1 population also with unrelated material of different leek types. Dosage calling was performed with the R package fitPoly (Voorrips *et al*., 2011; Zych *et al*., 2019)

### Linkage mapping

Linkage mapping in both F1 populations was performed using the R package polymapR (Bourke *et al*., 2018), according to the vignette of that package. The procedure consists of the following consecutive steps: filtering against markers and SNPs with too many missing data; merging duplicate F1 individuals; binning duplicate markers; assigning simplex x nulliplex (and analogous) markers to chromosomes and then to homologues for the two parents separately; matching the maternal and paternal chromosomes using simplex x simplex markers; assigning all other marker types to the simplex-nulliplex homologues; ordering the markers on the chromosomes; fine-tuning by successively eliminating ill-fitting markers. Finally, the duplicated markers that were set aside in the binning step were added back at the position of the marker representing the bin.

A consensus linkage map over the two populations was created by combining the pairwise linkage data (i.e., for each marker pair the recombination fraction and the LOD score) of the corresponding chromosomes from both populations. For marker pairs that occurred in the linkage data of both populations the combined recombination and LOD score were calculated by averaging the separate values, weighted by the squares of the LOD scores. With the combined linkage data marker ordering and fine-tuning was performed again.

Diagnostic plots were produced and a test for preferential pairing was performed, using functions from the polymapR package. Linkage maps were plotted using MapChart (Voorrips, 2002).

### Scaffolding

Selected contigs from the purging step were used as reference for mapping of the probe sequences of Axiom markers present on the final map. The probes were mapped using GMAP v2021-08-25 (Wu & Watanabe, 2005) with a large index, suitable for a genome larger than 232 Mb. They were then filtered to ensure a coverage of at least 0.98, an identity of at least 0.95, and only one mapping position. The mapping was combined with the original map and subsequently scaffolded using ALLMAPS v4 (Tang *et al*., 2015). Public reference genomes of *A. sativum*, *A. fistulosum* and *A. cepa* were obtained from NCBI RefSeq entries GCA_014155895.2, GCA_030737875.1, GCA_030737815.1. and GCA_030765085.1. Mapping of *A. porrum* markers against those assemblies was done similarly to the mapping described for scaffolding.

### Annotation

Repeats in the genome assembly were annotated using the RepeatModeler, RepeatClassifier and RepeatMasker tools using the combined RepBase (2014) and Dfam (2020) databases for classification of identified repeats (Bao *et al*., 2015; Hubley *et al*., 2016; Smith *et al*., 2013). Full length non-concatemer IsoSeq reads were mapped against the genome assembly using minimap2 *-ax splice -uf -- secondary=no -C5*. Braker v3 pipeline was then used for building gene models, gene prediction and gene annotation (Gabriel *et al*., 2024; Kovaka *et al*., 2019; Pertea & Pertea, 2020; Quinlan, 2014; Stanke *et al*., 2006, 2008). Efforts to reveal the centromeres, (sub-)telomeres and ribosomal DNA arrays were based on BLASTn searches using NCBI accessions MT374061.1, MT374062.1, MH017541.1 and MH017541.1 (Fu *et al*., 2019; Kirov *et al*., 2020). Ribosomal DNA arrays were annotated using Infernal v1.1.5 (Nawrocki & Eddy, 2013) and selected eukaryote 5S, 5.8S, 18S and 28S sequences from the RFAM database.

## Results

### *De novo* assembly

Sequencing of “Leidse prei 2018-94” DNA yielded 883 Gbp of PacBio HiFi data divided over 57.5 million reads (Supplementary Table S1, Supplementary Figure S2 & S3). In total, 4.2% and 5.0% of the hits were to plastid and mitochondrial databases respectively. Quality control showed an average read quality of Q30, which is generally considered a sufficient accuracy level for faithful genome reconstruction. K-mer analysis for k-mer size 21 revealed a GenomeScope profile showing the highest peak at a coverage of 14 and two additional minor peaks at 28 and 42 (Supplementary Figure S4), representing genomic sequences present in single copy or two and three copies, respectively. Modelling of the genome parameters such as genome size, heterozygosity and repetitiveness using the obtained distribution proved to be unsuccessful as the peaks did not diverge sufficiently, and only one broad peak was captured by the Genomescope2 model. K-mer analysis revealed strong AAAB and AB signals confirming the autotetraploid history of leek (Supplementary Figure S5). Rough estimation of genome size based on the total data set size and the first peak, under the assumption of full heterozygosity, was a 4n complement of 63 Gbp with 1n equaling to 15.75 Gbp.

*De novo* assembly resulted in a total assembly size of 70.7 Gbp consisting of 426,980 contigs with an N50 of 3.7 Mbp (Table 1). Primary and alternative assemblies sized to 38.9 and 31.9 Gbp with N50 sizes of 27.5 and 0.14 Mbp respectively. The largest contigs with an L50 index of 377 in the primary assembly, further supported the assignment of the primary assembly (Supplementary Figure S6). Contamination screening identified 49 and 2 hits to proteobacteria for primary and alternative assemblies respectively, which were removed. Additionally, 543 and 231 PacBio adaptor and primer sequences were detected in the primary and alternative contig sets and contigs were consequently either trimmed or broken. Single copy ortholog benchmarking, referencing the embryophyta lineage set, showed a very high completeness score of 97.2%, but also a high duplication rate of 95.2% as can be expected from a polyploid. For successful scaffolding with a linkage map, marker sequences must map to a single position in the genome and polyploid derived duplication must be minimized. To satisfy this criterion, we applied custom purging based on the BUSCO gene set, selecting 369 contigs (totaling 16.1 Gbp) while maintaining a high benchmark completeness score and reducing the duplication rate to 38.3%. While still a suboptimal percentage, this purging result combined with further selection on uniquely mapping markers would enable us to scaffold a pseudohaploid genome. The mapping rates of IsoSeq full-length transcripts against both the primary and purged contig sets were 96%, indicating that purging did not significantly impact transcriptome representation. This ensured that scaffolding and the final assembly retained high genomic integrity and transcript completeness.

**Table 1:** Assembly statistics for all subsequent stages of nuclear genome assembly and of chloroplast assembly.

| **Assembly:** | *Full* | *Primary* | *Alternative* | *BUSCO-purged* | *Scaffolded-all* | *Scaffolded-chromosomes* | *Chloroplast* |
| --- | --- | --- | --- | --- | --- | --- | --- |
| **Total size:** | 70,789,634,589 | 38,888,933,560 | 31,900,701,029 | 16,186,646,902 | 16,186,677,702 | 15,261,338,595 | 152,493 |
| **#Contigs:** | 426,980 | 59,335 | 367,645 | 369 | 61 | 8 | 1 |
| **N50:** | 3,760,608 | 27,546,430 | 142,303 | 56,398,703 | 1,910,501,443 | 1,910,501,443 | 152,493 |
| **L50:** | 2,020 | 376 | 35,492 | 96 | 3 | 3 | 1 |
| **GC-%:** | 35.64 | 35.61 | 35.67 | 35.56 | 35.56 | 35.56 | 36.73 |
| **#N:** | 0 | 0 | 0 | 0 | 30,800 | 30,800 | 0 |
| **Max contig size:** | 219,647,503 | 219,647,503 | 21,158,572 | 219,647,503 | 2,446,554,542 | 2,446,554,542 | 152,493 |
| **BUSCO-total:** | 1,614 | 1,614 | 1,614 | 1,614 | 1,614 | 1,614 | NA |
| **Complete:** | 1568 (97.2%) | 1560 (96.7%) | 1393 (86.3%) | 1560 (96.7%) | 1560 (96.7%) | 1521 (94.2%) | NA |
| **Single:** | 32 (2.0%) | 90 (5.6%) | 393 (24.3%) | 942 (58.4%) | 942 (58.4%) | 986 (61.1%) | NA |
| **Duplicated:** | 1536 (95.2%) | 1470 (91.1%) | 1000 (62.0%) | 618 (38.3%) | 618 (38.3%) | 535 (33.1%) | NA |
| **Fragmented:** | 7 (0.4%) | 12 (0.7%) | 4 (0.2%) | 12 (0.7%) | 12 (0.7%) | 9 (0.6%) | NA |
| **Missing:** | 39 (2.4%) | 42 (2.6%) | 217 (13.4%) | 42 (2.6%) | 42 (2.6%) | 84 (5.2%) | NA |
| **IsoSeq map rate:** | NA | 96% | NA | 96% | 96% | 96% | NA |

Assembly of the organellar genomes was successful only for the chloroplast but not for the mitochondrion. The reference-based chloroplast assembly has a total length of 152,493 bp and follows the conserved pattern of a long single copy section (LSC) and short single copy section (SSC) interspersed by an inverted repeat. This result indicates that the chloroplast genome is highly conserved and structurally resembles known chloroplast reference genomes. This allowed for successful assembly through reference-based methods, whereas the failure to assemble the mitochondrial genome may suggest greater complexity or divergence in its structure, possibly requiring more specialized assembly approaches or higher-quality data.

### Linkage mapping

The linkage maps of both F1 populations, and the integrated map, consisted of 8 linkage groups, corresponding to the eight chromosomes of leek. Overall statistics of the maps are shown in Table 2. Diagnostic plots for the integrated map are shown in Supplementary Figure S7, and scatterplots of the Population 1 and Population 2 maps versus the integrated map in Supplementary Figure S8. The integrated map and the F1 population 1 map, the latter having the target “Leidse prei 2018-94” plant as father, were quite similar, although the integrated map contains more markers. The length of the chromosomes in both maps did not differ much. The map of F1 Population 2 showed some significant differences with the integrated map and the Population 1 map. The map of Population 2 was much sparser, and chromosomes 6 and 8 in this map were much shorter than the other maps. Also, the lengths of chromosomes 3 and 7 were larger than those of the other two maps, containing larger gaps. These map features point to a lower quality for the map constructed from the second population. The integrated map of chromosome 6 is based only on the linkage data of F1 population 1; it is identical to it, except that a few markers that were added in Population 2 showed an identical segregation as the markers present in the Population 1 map.

**Table 2:** Linkage map statistics.

| Chromosome | Integrated map |  | Population 1 |  | Population 2 |  |
| --- | --- | --- | --- | --- | --- | --- |
|  | markers | length (cM) | markers | length (cM) | markers | length (cM) |
| 1 | 1927 | 137.1 | 1463 | 136.8 | 773 | 135.8 |
| 2 | 2273 | 146.3 | 1593 | 142.2 | 1133 | 147.6 |
| 3 | 1825 | 138.5 | 1373 | 128.4 | 683 | 152.9 |
| 4 | 1235 | 150.9 | 755 | 146.7 | 611 | 150.0 |
| 5 | 1264 | 138.3 | 768 | 136.9 | 662 | 138.6 |
| 6 | 1107 | 138.5 | 1090 | 138.5 | 557 | 99.0 |
| 7 | 1115 | 133.2 | 833 | 123.5 | 445 | 153.0 |
| 8 | 683 | 139.8 | 587 | 143.1 | 124 | 63.7 |

The markers on the integrated linkage map were mostly concentrated in dense clusters near the ends of all chromosomes (Figure 1). A similar distribution was observed in the separate maps for the first and second F1 populations. However, in the map of the second population, the left distal regions of chromosomes 6 and 8 were missing (not shown). Since the markers were derived from expressed sequences, this suggests that a large part of the expressed genes are in blocks with low recombination frequency. In Population 1 the occurrence of double reduction was studied based on simplex x nulliplex markers of both parents. At both telomeres of all chromosomes a level of 4%-5% double reduction was observed, indicating the occurrence of quadrivalents and polysomic inheritance.

**Figure 1:**
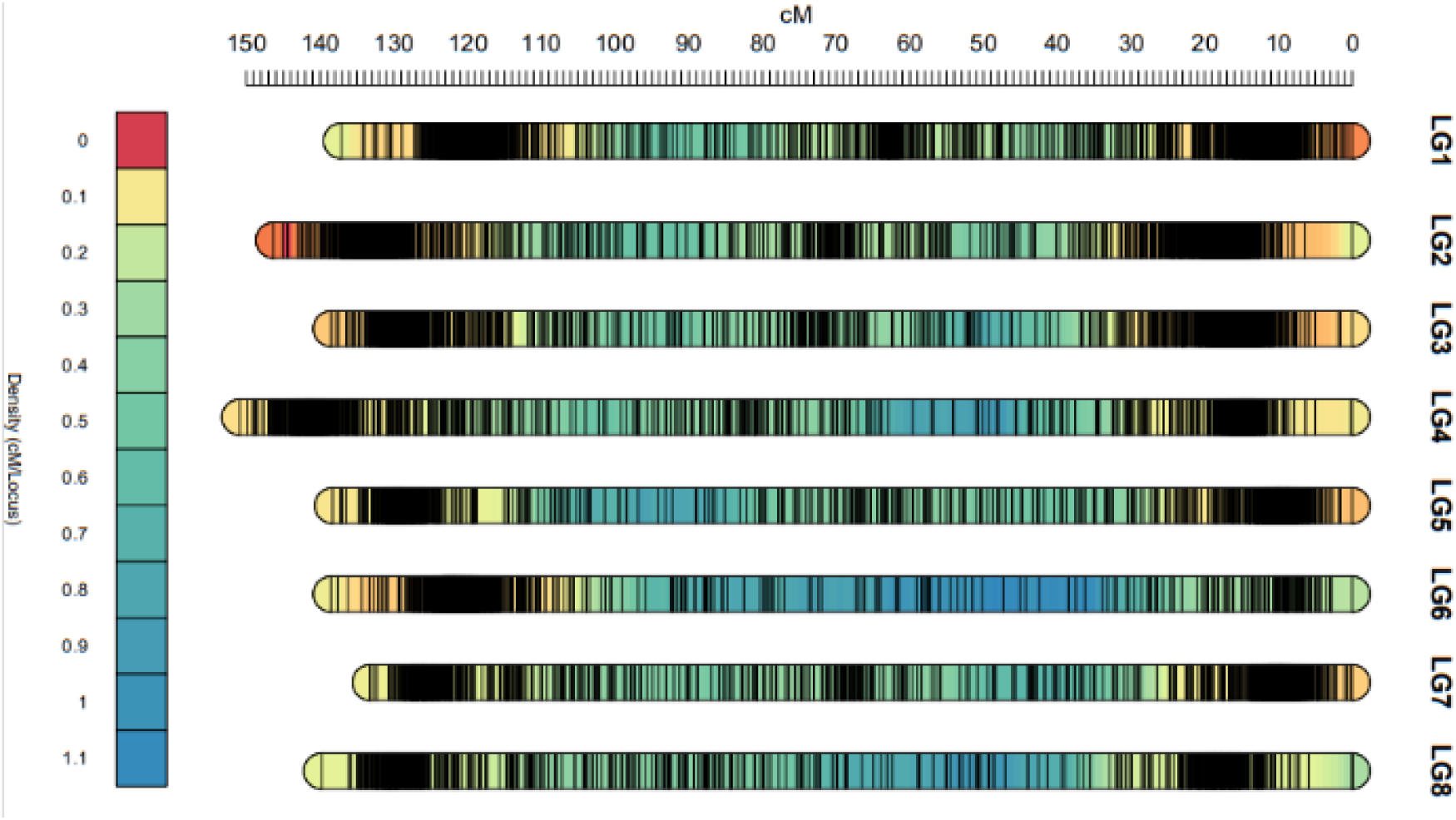
Marker density in cM/locus for the eight linkage groups established in the integrated genetic map of Allium porrum. Each vertical stripe represents one marker. Highly dense regions tend to color black and generally can be seen at both distal ends and center of each linkage group.

### Scaffolding

Mapping of marker sequences to 369 selected contigs, representing a minimal set with maximum BUSCO completeness, showed that 11,143 out of 11,429 markers (97.5%) were successfully mapped. Further filtering based on alignment coverage, identity, and mapping ambiguity removed 835, 300, and 3,123 markers, respectively. The remaining 7,184 mapped markers were used to orientate and order the 369 contigs with the linkage map as template to construct a chromosome-level genome assembly. Of the 316 contigs anchored with the map, 303 could be oriented using at least two aligned markers. The placed contigs represented 94.3% of the total sequence length, with 15.9 Gbp of the 16.2 Gbp scaffolding input mapped to the linkage map. Of this, 15.3 Gbp was anchored to chromosomes, and 15.0 Gbp was also oriented. Unplaced sequences included 25 contigs lacking marker information and 28 contigs were omitted due to unresolved conflicts with the linkage map. The largest chromosome, chromosome 2, measured 2.45 Gbp, while the smallest, chromosome 8, was 1.40 Gbp. Overall, the correlation between genetic and physical positions was high, with Spearman’s rank correlation coefficient (ρ) ranging from 0.96 for chromosome 4 to 0.66 for chromosome 6, with most chromosomes exceeding 0.9 (Supplementary Figure S9).

### Repeat and gene annotation

*De novo* repeat modeling and annotation, using RepBase and curated DFAM databases, identified 4,312 repeat models. The largest group remained unclassified, comprising 1,725 unknown repeat families and 1,688 unknown LTR retrotransposon families. Among the classified repeats, 391 belonged to the LTR/Copia family, while 361 were classified as LTR/Ty3 (formerly known as Gypsy). Overall, repeat masking covered 81.51% of the leek genome assembly, highlighting its highly repetitive nature, dominated by both known and unknown transposable elements. Gene annotation of the repeat masked genome resulted in 66,021 genes of with an average length of 6,553 bp and a mean 3.7 exons per gene. The repeat and gene distribution can be described as uniformly distributed along the physical chromosomes and no clear evidence was found of centromere and telomere associated sequences (Supplementary Figure S10).

### Scaffolded genome reveals distinct recombination landscape

The Marey maps (Figure 2) show a comparison between the generated linkage map and the physical map that was obtained by *de novo* assembly. Overall, the maps indicate a sufficient coverage of markers along the chromosomes. The densely populated areas on the linkage map make up the chromosome ends. Because the array design was based on transcriptome derived variants, the marker distribution shows the density of genic regions which are present quite uniformly along the physical chromosomes in leek (with slightly higher densities in distal parts of the linkage map). For all chromosomes a full sigmoid like pattern can be observed, with the exception of chromosome 6. This chromosome shows a partial sigmoid like pattern with a flat part extending the chromosome end. Most of the chromosomes lack markers in the middle section, possibly representing the centromere region. The flatter parts at the chromosome ends indicate a smaller distance between successive markers on the genetic map, while the middle parts of each chromosome show relatively larger genetic distances between successive markers. This pattern thus points to a relatively high recombination frequency in the middle of the chromosome compared to the chromosome ends. Apparently, recombination in *A. porrum* occurs mainly in the regions directly adjacent to the centromere and not in the distal chromosome ends, consistent with the cytological observation that chiasmata in *A. ampeloprasum* are predominantly localized proximally to the centromere (Kollmann, 1972).

**Figure 2:**
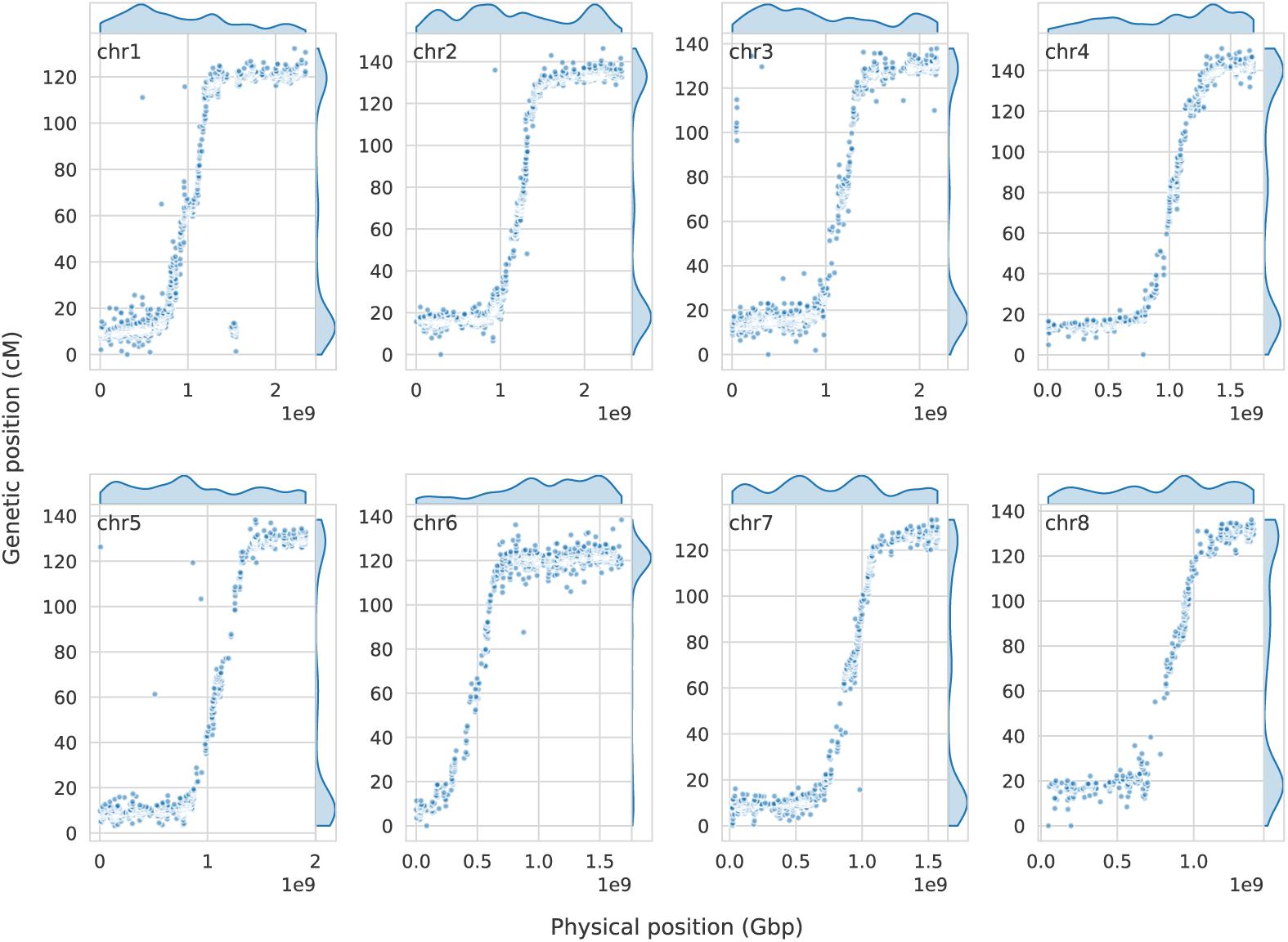
Marey maps displaying physical (x-axis) and genetic positions (y-axis) of markers. Density estimates are displayed along the axes for each chromosome. The sigmoid-like patterns reveal the proximal recombination landscape in A. porrum. Density estimates show a large number of markers in regions with minimal recombination on the genetic map.

### Genome comparison to *A. fistulosum*, *A. sativum and A. cepa*

A comparison of physical marker positions from our genetic map with physical positions on other recently published high-quality genome assemblies provides insight into large structural chromosome rearrangements. Mapping the marker sequences from the final genetic map to *Allium fistulosum* (Welsh onion/bunching onion) (Liao *et al*., 2022; Hao *et al*., 2023) yielded 1,135 unambiguous mapping positions. As shown in Figure 3, large intrachromosomal inversions are present on chromosomes 1, 2, 3, 5, and 6. Although a few markers suggest possible translocations by mapping to different chromosomes—indicated by deviations in the color code displayed in the chromosome synteny panels of Figure 3—no major interchromosomal rearrangements were observed between leek and bunching onion (*A. fistulosum*).

**Figure 3:**
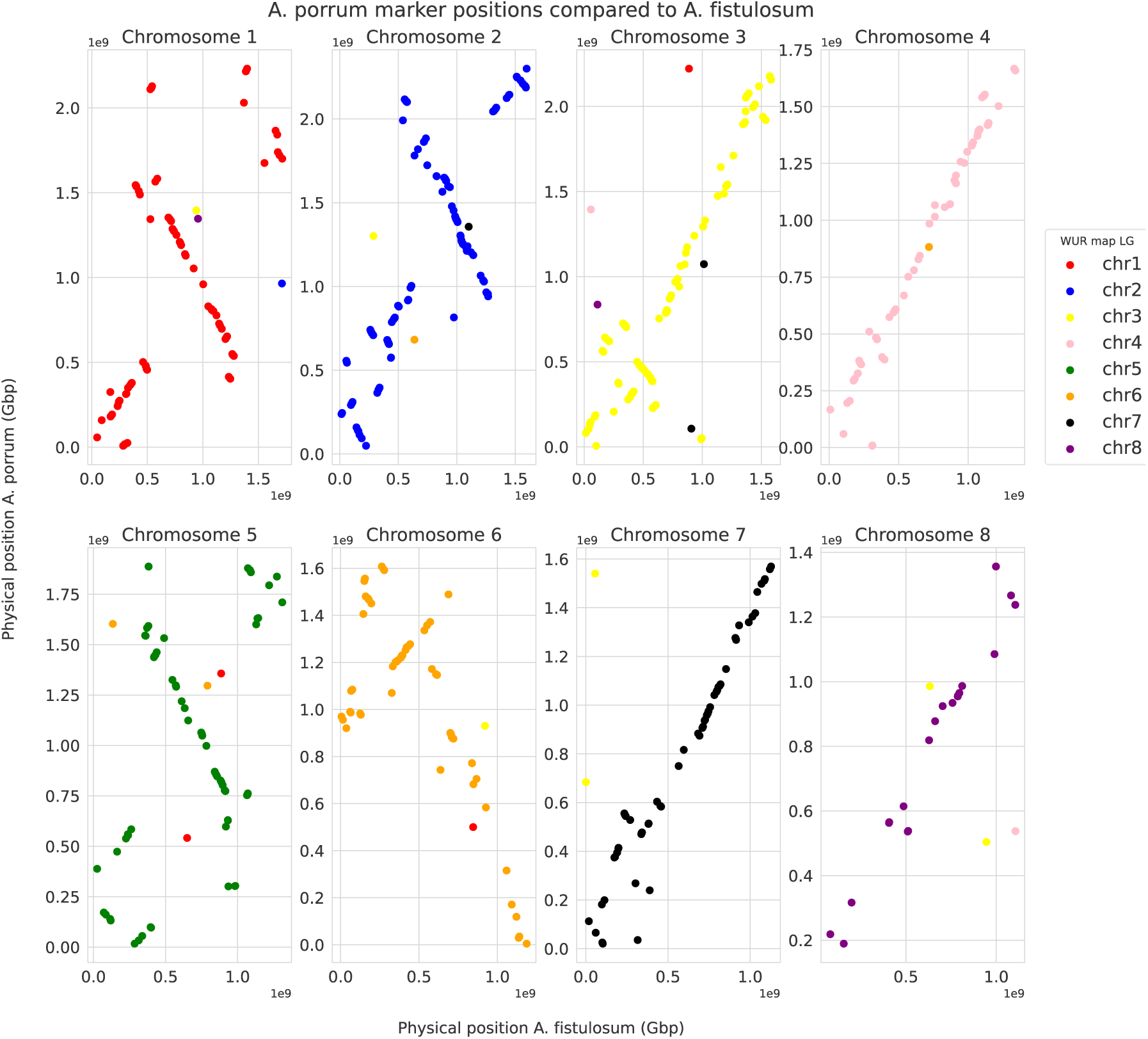
Mapping of markers on A. porrum (y-axis) and A. fistulosum (x-axis) reveal several large structural variations such as inversions of 1 Gbp on chromosomes 1, 2 and 5. Chromosomes 4 and 7 seem to be overall more colinear at this resolution.

Furthermore, a comparison between the final genetic map and *A. sativum* (Sun *et al*., 2020) identified 4,785 mappable marker positions, revealing synteny between leek chromosomes 1, 2, 3, 6, and 8 with *A. sativum* chromosomes 6, 3, 8, 4, and 1, respectively. However, these chromosomes also exhibit synteny breaks due to intrachromosomal inversions (Supplementary Figure S11). Additionally, large interchromosomal rearrangements were observed for leek chromosomes 4, 5, and 6. Specifically, leek chromosome 2 was partially syntenic with *A. sativum* chromosomes 2 and 7, leek chromosome 5 showed partial synteny with *A. sativum* chromosomes 2 and 5, and leek chromosome 6 was partially syntenic with *A. sativum* chromosomes 5 and 7. However, a comparison with *A. sativum* from Hao and coworkers (2022) identified 5,013 shared markers but did not reveal intrachromosomal rearrangements (Supplementary Figure SX). This discrepancy may reflect differences in genome assembly quality, as suggested by Hao *et al*. (2022). Alternatively, it is possible that these differences result from accession-specific rearrangements in *A. sativum*. A comparison with *A. cepa* (Hao *et al*., 2022) identified 1,116 shared mappable markers and revealed synteny breaks due to intrachromosomal rearrangements across all chromosomes (Supplementary Figure S11). In contrast, no major interchromosomal rearrangements were observed between leek and onion. Considering the higher number of shared markers and the smaller size of inversions, *A. ampeloprasum* appears to be more closely related to *A. sativum* than to *A. cepa* or *A. fistulosum*. This result is consistent with the phylogenetic relationships among *Allium* species as previously reported by Hirschegger *et al*. (2010).

## Discussion

### First chromosome-scale reference genome of *A. porrum*

The first genome sequence of a highly heterozygous tetraploid *Allium porrum* is a valuable addition to the collection of Allium genomes that have currently been reconstructed, as the current collection does not contain such a complex genome yet. The total size of the 15.26 Gbp anchored genome is slightly smaller than the recently published genomes of onion (15.78 Gbp) and garlic (15.52 Gbp), but bigger than the Welsh onion genome of 10.48 Gbp. Compared to the N50 contig size of 81.66, 109.82 and 507.27 Mbp for onion, garlic and Welsh onion respectively, the N50 leek contig size of 57.37 Mbp is notably smaller. This is partly due to the relatively low coverage per haplotype combined with the high heterozygosity and possibly a smaller sequence library insert size. Since the Allium genomes reported by Hao *et al.,* 2023 were also constructed by HiFi based technology, we regard the difference in N50 contig size less likely to be the result of the sequencing technology used. Several findings have reported on genome size for *A. ampeloprasum*. Our leek genome assembly size significantly exceeds the flow cytometric values reported by Arumuganathan & Earle (1991), who found a 2C value of 50.27 pg for *A. ampeloprasum*, corresponding to a single-copy genome size of 12.29 Gbp. Ricroch *et al*. (2005) reported that the genome size of tetraploid *A. porrum* is 50.7 ± 0.7 pg, corresponding to *n*=12.45 Gbp, and additionally noted that the diploid *A. ampeloprasum* has a haploid genome size of 16.37 Gbp. Ohri *et al*. (1996) summarized findings on tetraploid *A. porrum*, which range from 11.78 Gbp to 31.93 Gbp for a single genome copy. Overall, the size of the leek genome assembly we report here falls within the previously reported genome size ranges for both *A. ampeloprasum* and *A. porrum*.

The use of PacBio HiFi reads with an average read quality score of Q30 was a key factor contributing to the high quality of the reconstructed leek genome. The high quality of the presented assembly is substantiated by the high completeness of conserved single copy ortholog (BUSCO) genes. The 97.2% completeness slightly exceeds the benchmark results for onion (96.4%), garlic (92.6%) and Welsh onion (96.6%) genomes. Our annotation efforts show that the number of 66k genes in leek is in the same range as found by *Hao* in garlic and onion, which are similar sized genomes. Annotation of repetitive sequences is different however, as the percentage of the genome marked as repetitive is around 11-13 percentage points lower. Marker-based scaffolding with probe sequences designed from transcriptome data and BUSCO-based purging are possible causes for the exclusion in the scaffolding process of contigs consisting of mainly repeat sequence. This depletion would lower the total repeat sequence relatively to the total assembly size. In addition to the high BUSCO scores, the high collinearity with an integrated map from two mapping populations confirms that the order of markers on the generated contigs aligns precisely with linkage calculations. Furthermore, the successful anchoring and orientation of a significant proportion (94.3%) of the genome assembly onto the integrated linkage map, along with the high correlation between genetic and physical positions, demonstrates the robustness and accuracy of the assembly process. The few unresolved contigs and lower correlation on chromosome 6 suggest areas for further refinement, but overall, the results provide a highly contiguous and well-ordered chromosome-level assembly that will support downstream genomic analyses.

### Marker-rich genic regions in distal clusters on linkage maps and recombination predominantly occurring near centromeres

Understanding gene distribution and recombination behavior across a wide range of Allium genomes is crucial for developing effective breeding programs. Furthermore, insight into marker distribution can aid in more efficient breeding by introducing marker-assisted selection. By leveraging dense markers in gene-rich areas, breeders can more accurately track desired traits, improving the speed and efficiency of developing new leek varieties. The markers we developed from mapped transcript sequences primarily target expressed leek genes. Surprisingly, we observed a pronounced clustering of markers towards the chromosome ends on the linkage map, whereas recombination predominantly occurred proximal to the leek centromere (Figure 2). This clustering of markers is partly the result of a relatively low recombination frequency towards the ends of the leek chromosomes, as we observed a relatively more even marker distribution on the physical map (Figure 2). Nonetheless, this contrast suggests that while breeders may focus on gene-rich regions for selecting desirable traits, recombination in the distal leek chromosome regions is limited, potentially constraining reshuffling of traits to new generations. Consequently, understanding the genetic landscape of *Allium* species is essential for optimizing breeding approaches.

Notably, variations in recombination behavior have been observed across different *Allium* species and subspecies. In tetraploid *A. porrum*, Levan (1940) reported a predominantly proximal localization of chiasmata, being cytological manifestations of crossovers. Further cytological studies by Kollmann (1972) on *A. ampeloprasum* subspecies (*ampeloprasum* and *truncatum*) revealed distinct recombination patterns: while *A. ampeloprasum* exhibited chiasmata localized near the centromere, *A. truncatum* displayed subterminal or terminal localization. Interestingly, within each subspecies, chiasma localization appeared independent of ploidy level. In addition, pollen abortion analyses revealed pollen viability to be the most stable in tetraploids and less for other ploidy levels for both subspecies. In addition, the predominant proximal localization of chiasmata seems to reduce multivalent chromosome formation in 4x, 5x, and 6x plants of subsp. *ampeloprasum*. However, no results pointing to a correlation between chiasma localization and pollen viability were reported (Kollmann, 1972). Profound differences in recombination behavior have also been observed between diploid Allium species. In onion (*A. cepa*), crossovers predominantly occur in the distal chromosome regions, whereas in *A. fistulosum*, they are mainly localized proximally (Emsweller and Jones, 1935a; Kudryavtseva *et al*., 2023). Moreover, chiasmata in an *A. cepa* × *A. fistulosum* hybrid were found to shift significantly toward the distal regions of *A. fistulosum* homologous chromosomes, suggesting genetic control over crossover localization. This observation supports the hypothesis of Emsweller & Jones (1935b) that *A. cepa* and *A. fistulosum* possess dominant and recessive genes, respectively, regulating distal and interstitial chiasmata localization. Their study furthermore showed that progeny from backcrosses of *cepa* x *fistulosum* hybrids to *cepa* and *fistulosum* exhibited pronounced differences in blooming and fertility. However, no correlation was found between fertility and percentage of good pollen. Interestingly, results suggest a correlation between fertility and chiasma localization as the most fertile plants displayed interstitial chiasmata, while sterile plants displayed terminal chiasmata. However, any conclusion from these observations should be drawn with caution, as the number of plants studied was rather limited (Emsweller & Jones, 1935b). Nonetheless, these recombination differences across Allium species may have significant implications for breeding and warrant further investigation into Allium meiotic recombination landscapes.

A comparison of our genetic and physical chromosome maps of *A. porrum* (Figure 2) further supports the previously reported findings and indicate that recombination occurs predominantly in the central parts of leek chromosomes, containing functional centromeres as demonstrated previously by Levan (1940). Subsequent FISH analysis in *A. cepa* and *A. fistulosum* showed the co-localization of repetitive centromeric sequences like those found in many plant species (Kirov *et al*., 2020), suggesting that recombination in leek also occurs mostly proximally to the centromeres. We were however unable to obtain any conclusive results on the inclusion of centromeres in our assembled genome, presumed to be caused by the absence of markers in the centromere itself. In this study, we combined information on recombination frequencies between markers obtained through linkage mapping with their physical genome sequence positions, further confirming the proximal localization of recombination in leek chromosomes and further substantiating the existence of a dichotomy in recombination behavior in *Allium*. To our knowledge, this is the first such comparison in the *Allium* genus.

## Abbreviations

LOD: log of odds
SNP: single nucleotide polymorphism
HiFi: High fidelity
HMW: high molecular weight
CO: Crossover

## Supplementary materials

Tables:

| Number: | Title: | Description: |
| --- | --- | --- |
| S1 | HiFi sequencing overview | Contains an overview of per SMRTcell stats on data production |
| S2 | Integrated genetic map | The linkage map generated from two populations merged (average) with the exception of chromosome 6 |
| S3 | Linkage map stats per LG | Overview of statistics for each linkage group |
| S4 | Annotation overview | Overview of annotation statistics per chromosome |

Figures:

| Number: | Title: | Description: |
| --- | --- | --- |
| S1 | Custom purging pseudocode | Describes method on how purging of primary contigs was done |
| S2 | HiFi read lengths | Read length distribution of all combined HiFi reads |
| S3 | HiFi mean read quality | Mean read quality distribution of all combined HiFi reads |
| S4 | Genomescope2 output | K-mer histogram and genomescope2 model fit for k = 21 |
| S5 | Smudgeplot | Smudgeplot for detection of ploidy and ploidization events |
| S6 | L50 plots | Plots showing the contiguity increase for each assembly iteration |
| S7 | Linkage map QC | For each linkage group there are pairwise comparisons of estimated recombination frequency and observed recombination frequencies and LOD scores |
| S8 | Linkage map comparison | Comparison of the separate maps to the integrated map |
| S9 | Scaffolding per linkage group | Shows how each chromosome is built from contigs and the marker order from AllMaps output |
| S10 | Feature density on chromosome level assembly | Visualized the feature density for repeats, genes, markers, ribosomal DNA and some satellite repeats |
| S11 | Structural homology to *A. sativum* and *A. cepa* | Genome comparisons based on marker mappings from our integrated map to two different version of the garlic genome and the onion genome |

## Acknowledgements

This project was supported by the Netherlands Top Consortium for Knowledge and Innovation (TKI project TU-18013). We also thank ENZA Zaden Research & Development B.V., Bejo Zaden B.V., Hazera Seeds B.V., and Nunhems Netherlands B.V. for their support.

## Data availability

The genomic and transcriptomic data used in this study are available under NCBI BioProject accession PRJNA1231124.

