## Supplementary Figures for "High-resolution genome and genetic map of tetraploid *Allium porrum* expose pericentromeric recombination"

#### S1 BUSCO-based purging pseudocode

```
C=[contigs] #Sorted by contig length in descending order
buscos_in_selected = []
selected_contigs = []
For contig in C:
        If any busco_on_contig not in buscos_in_selected
                append busco_gene to buscos_in_selected
                append contig to selected contigs
        else:
                next
```

#### S2 HiFi read lengths

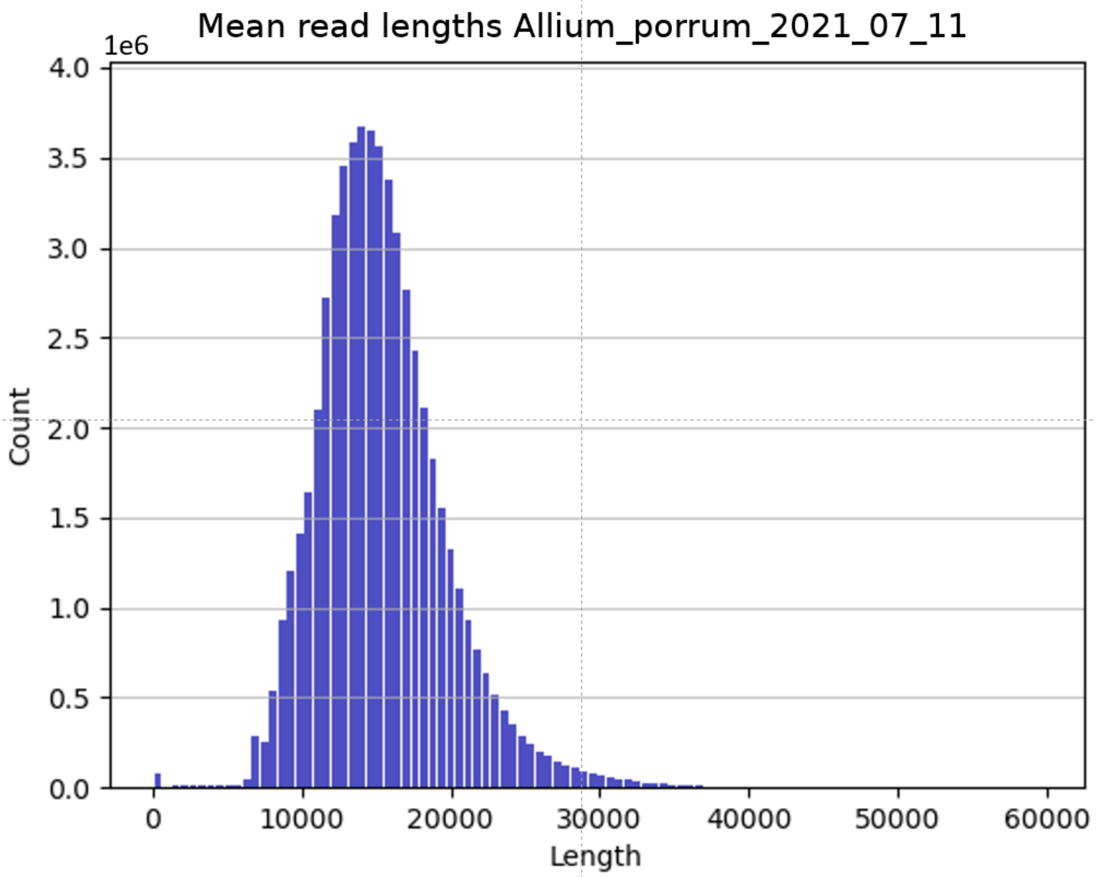

S3 HiFi mean read quality

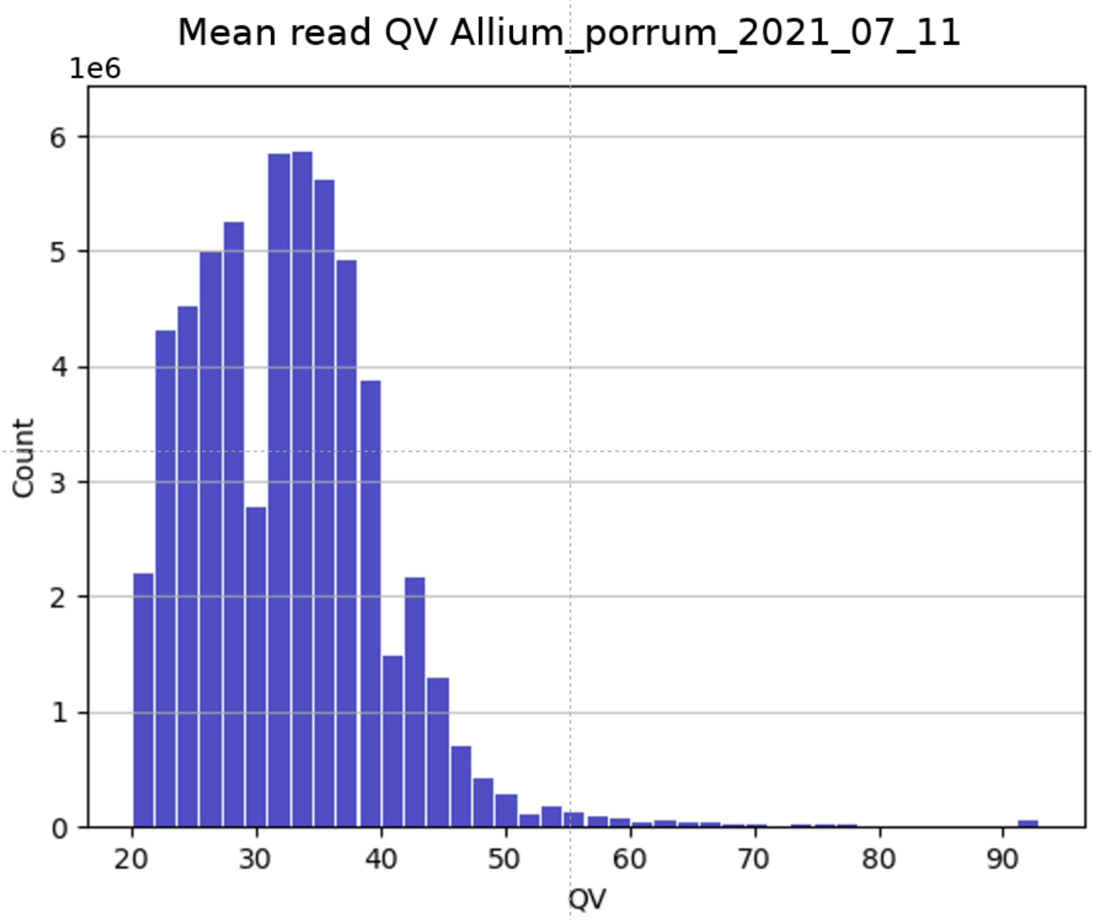

#### S4 GenomeScope2 output

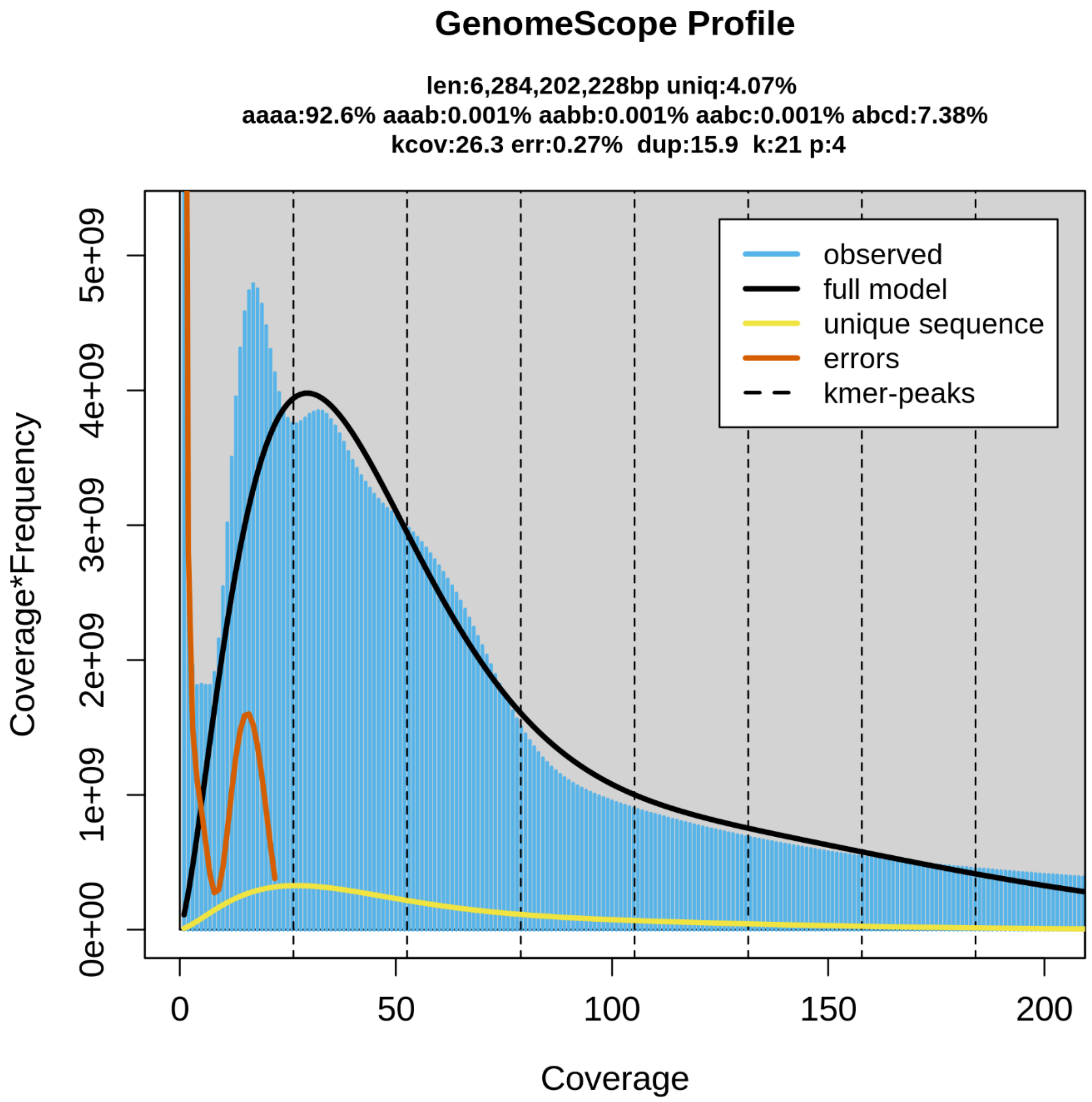

#### S5 Smudgeplot

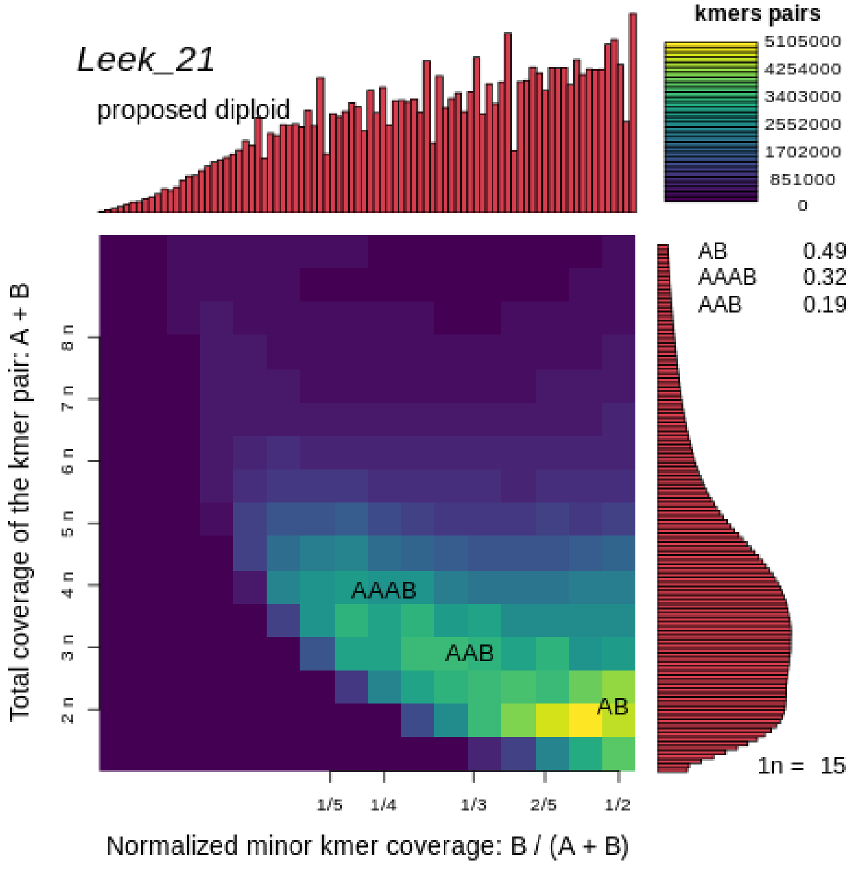


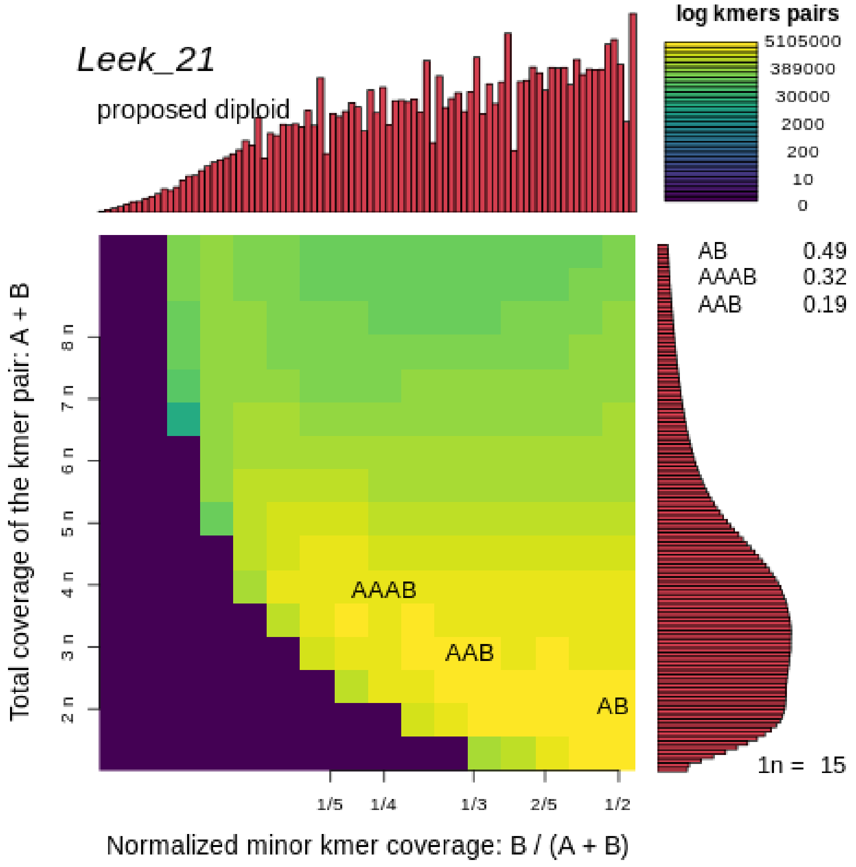

### S6 L50 plots

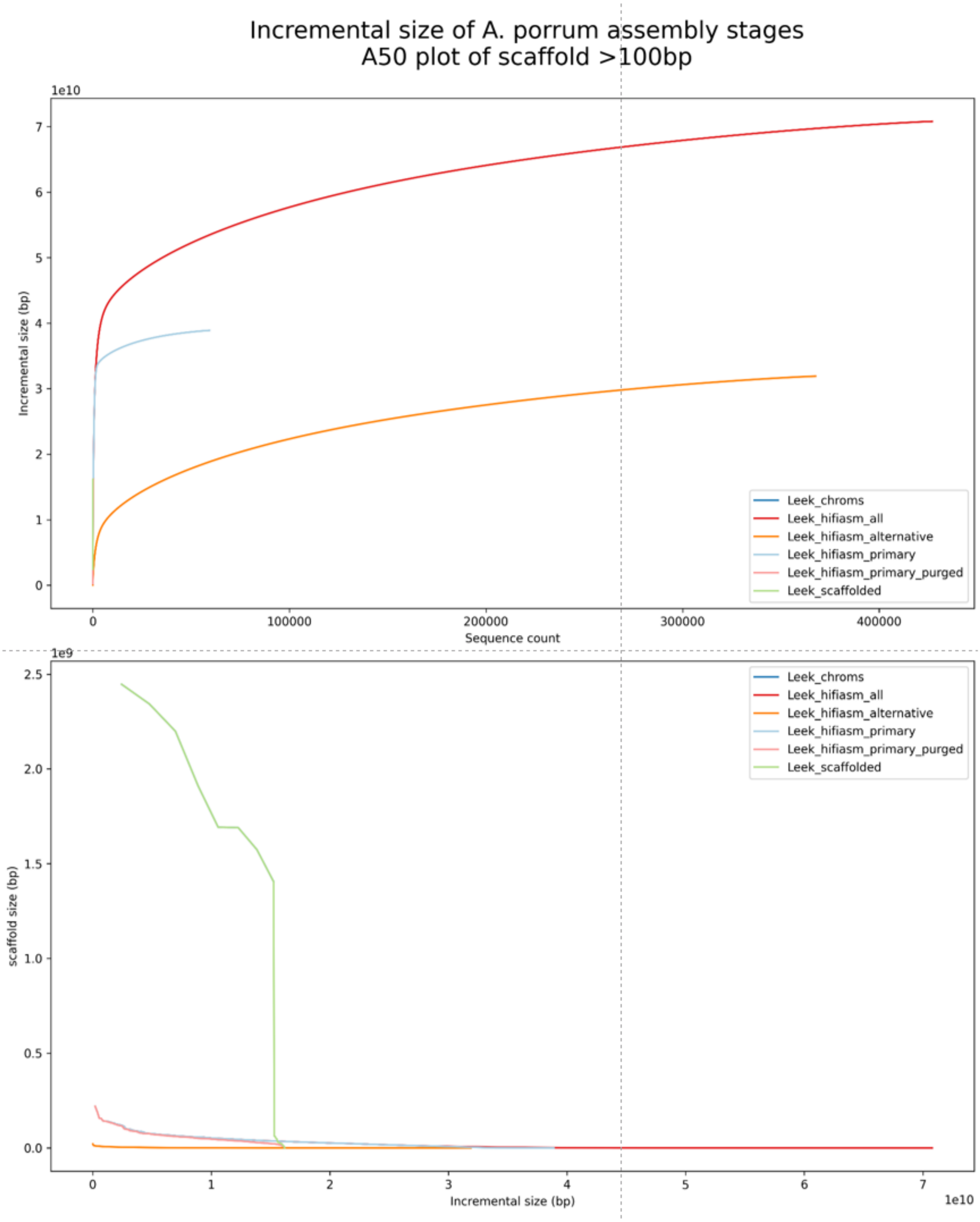

Incremental size of A. porrum assembly stages
A50 plot of scaffold >100bp
1e10
Incremental size (bp)
Sequence count
0
100000
200000
300000
400000
Leek_chroms
Leek_hifiasm_all
Leek_hifiasm_alternative
Leek_hifiasm_primary
Leek_hifiasm_primary_purged
Leek_scaffolded
1e9
scaffold size (bp)
Incremental size (bp)
1e10

#### S7 Linkage map QC

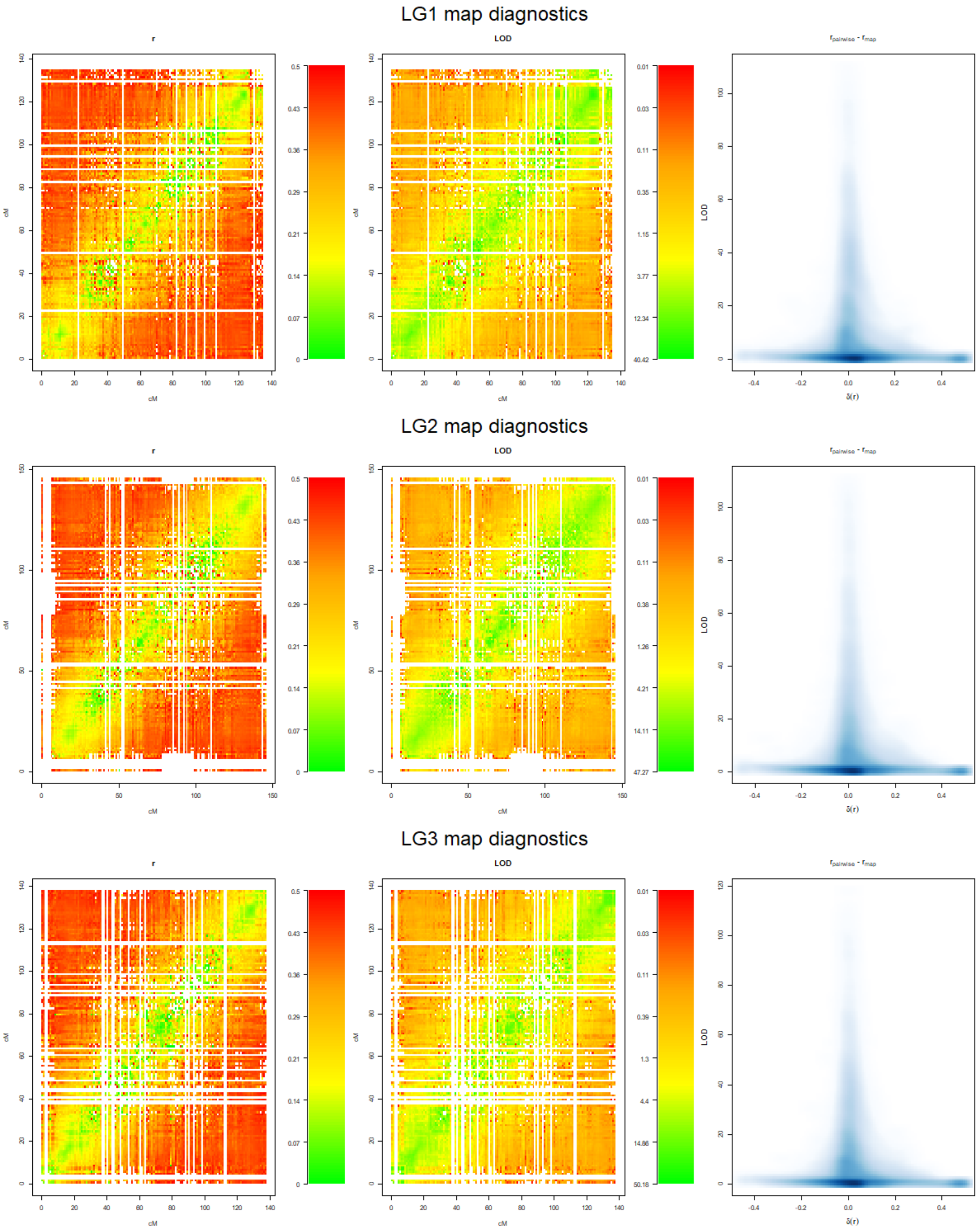

LG4 map diagnostics

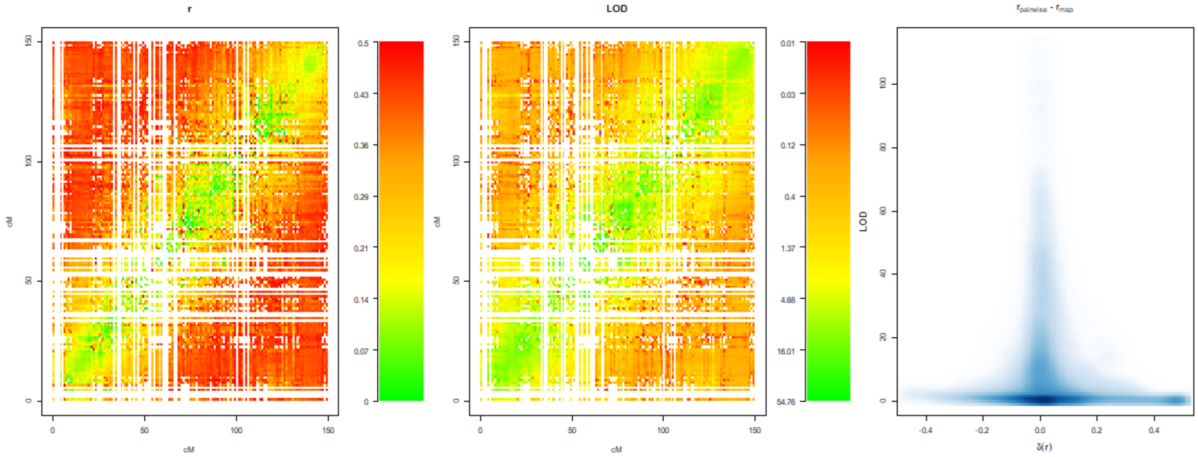


LG5 map diagnostics

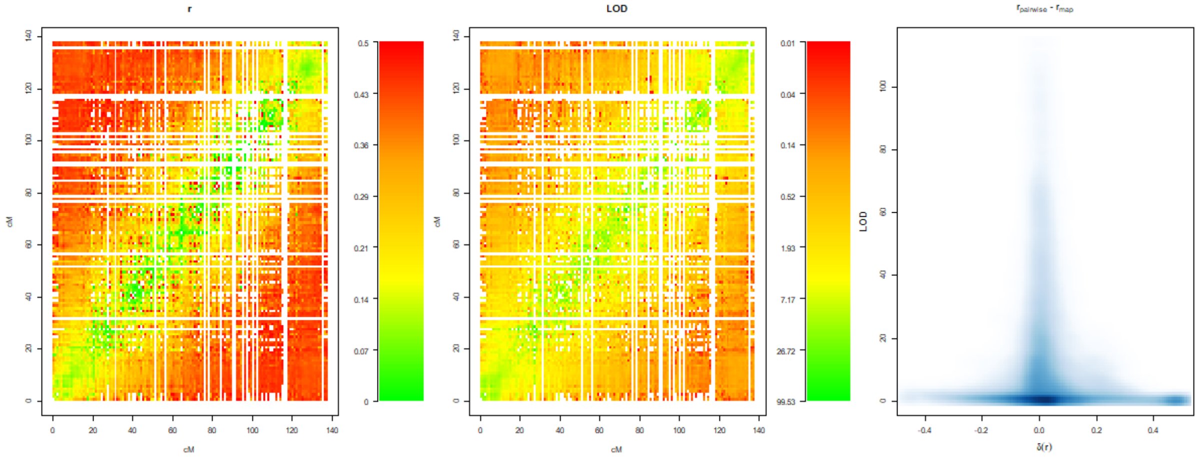


LG6 map diagnostics

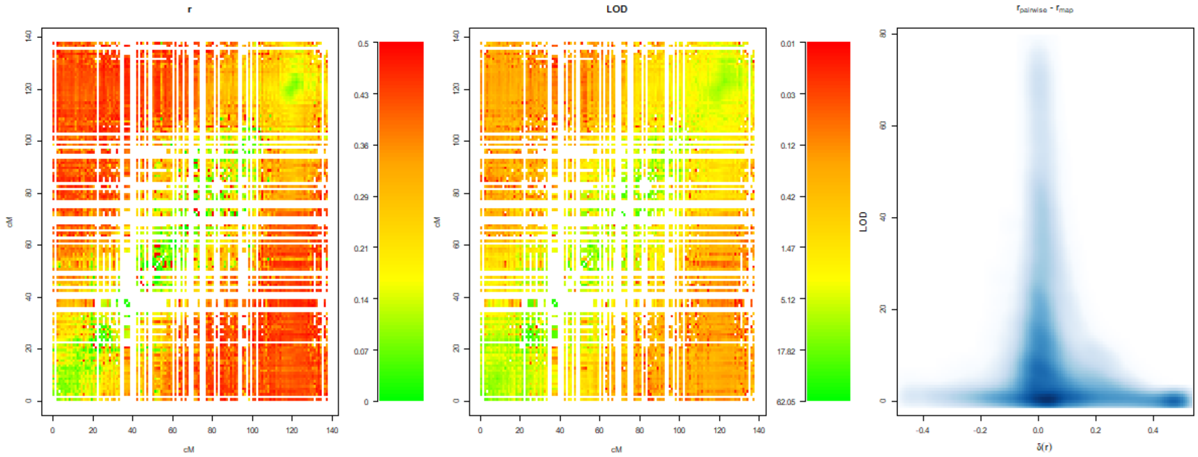

LG7 map diagnostics

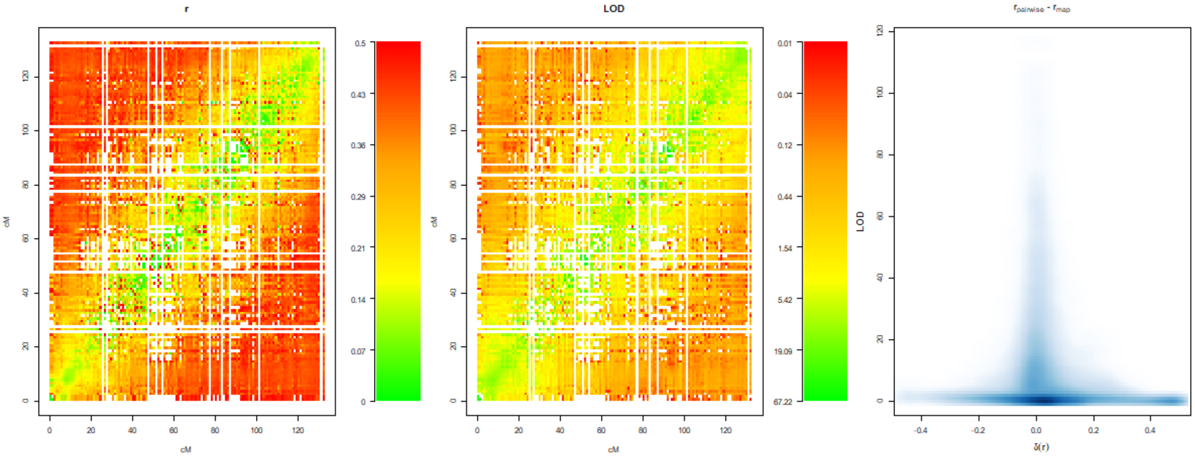


LG8 map diagnostics

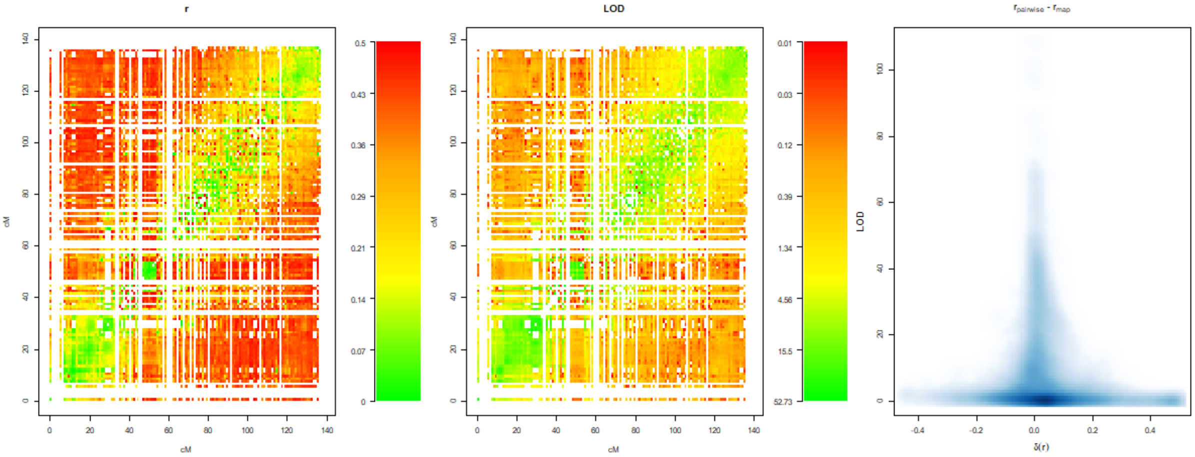

#### S8 Linkage map comparison

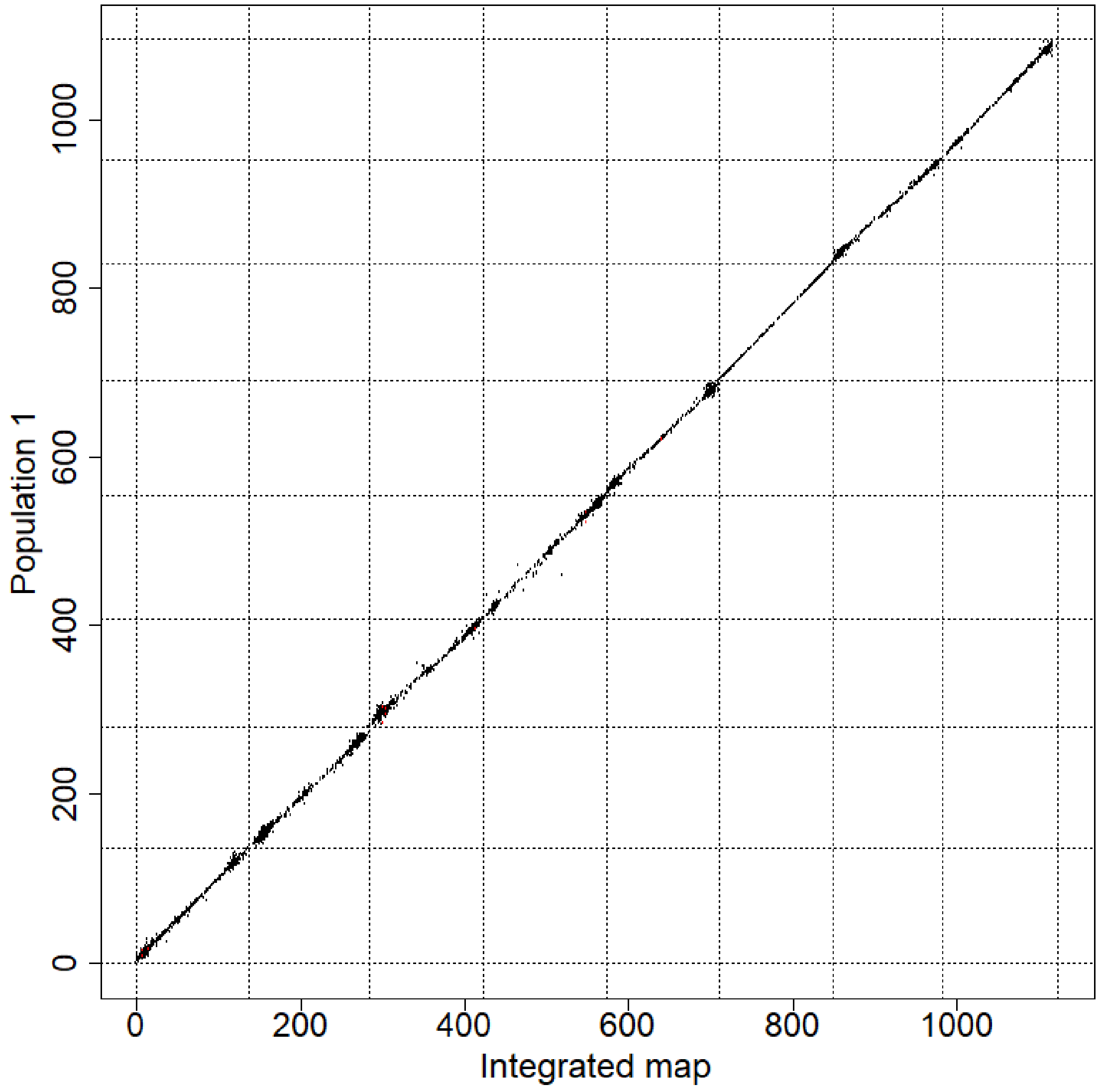

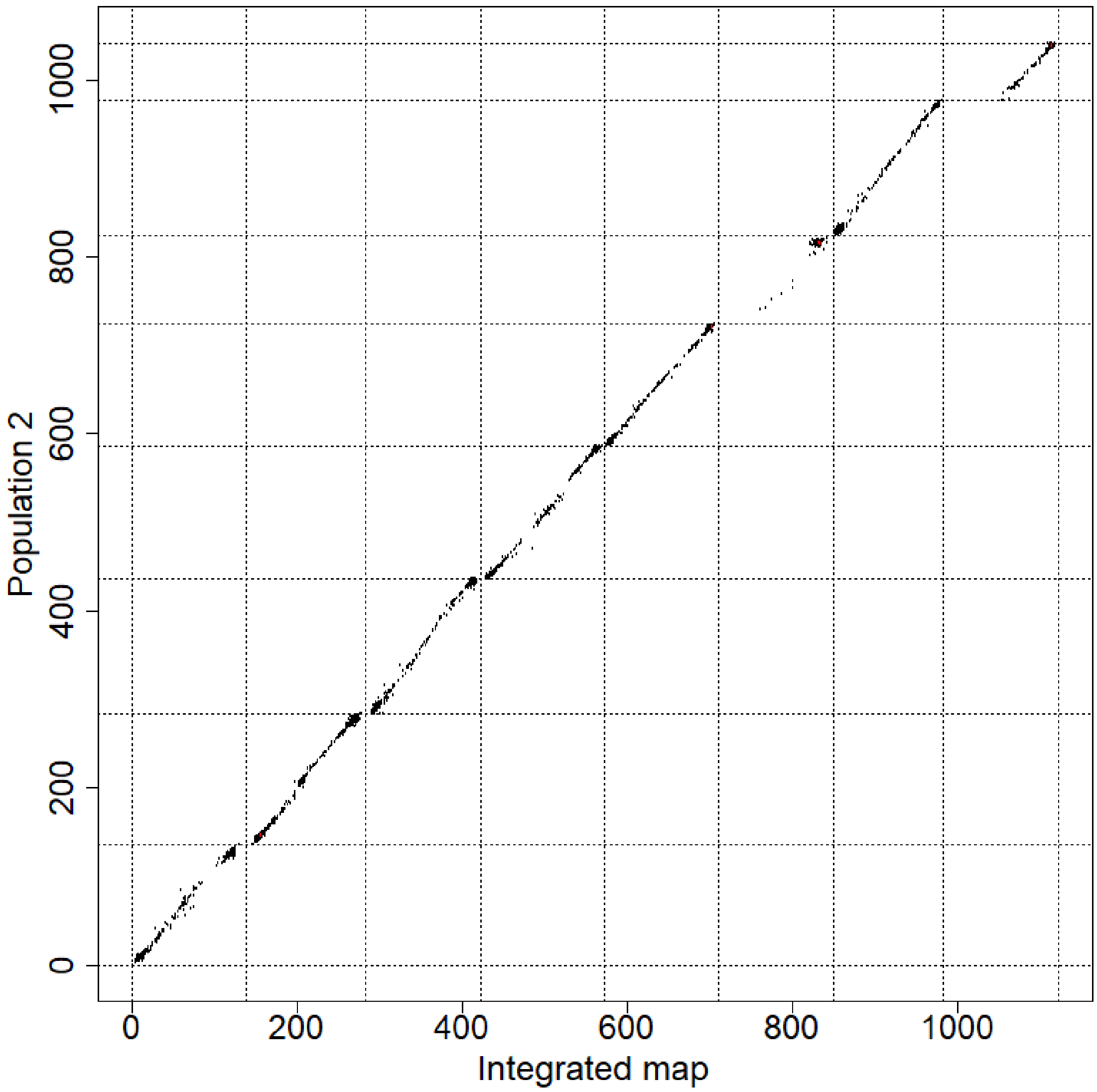

Population 2
0
200
400
600
800
1000
Integrated map

#### S9 Scaffolding per linkage group

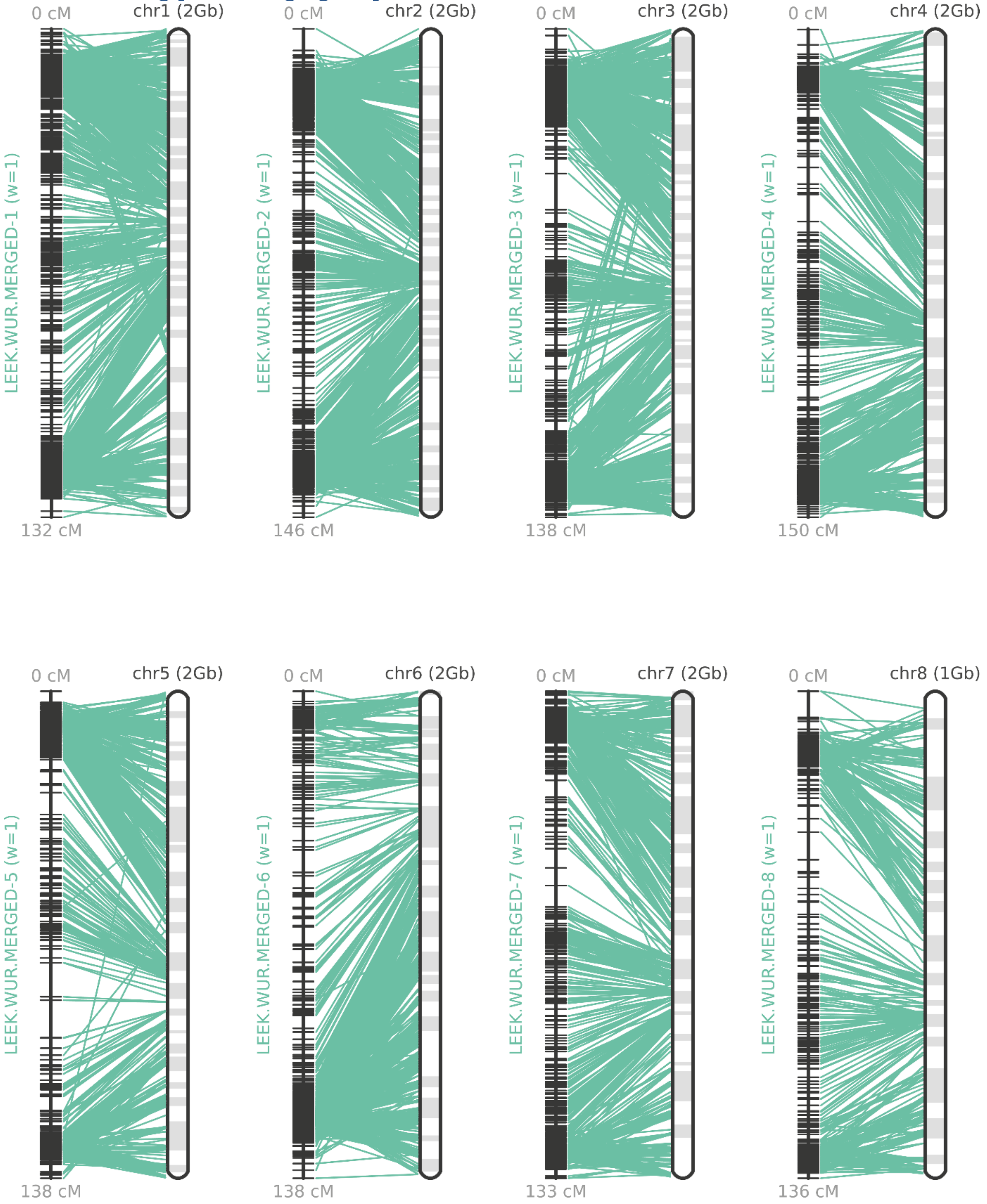

#### S10 Marker density, rDNA annotation, centromere localization markers

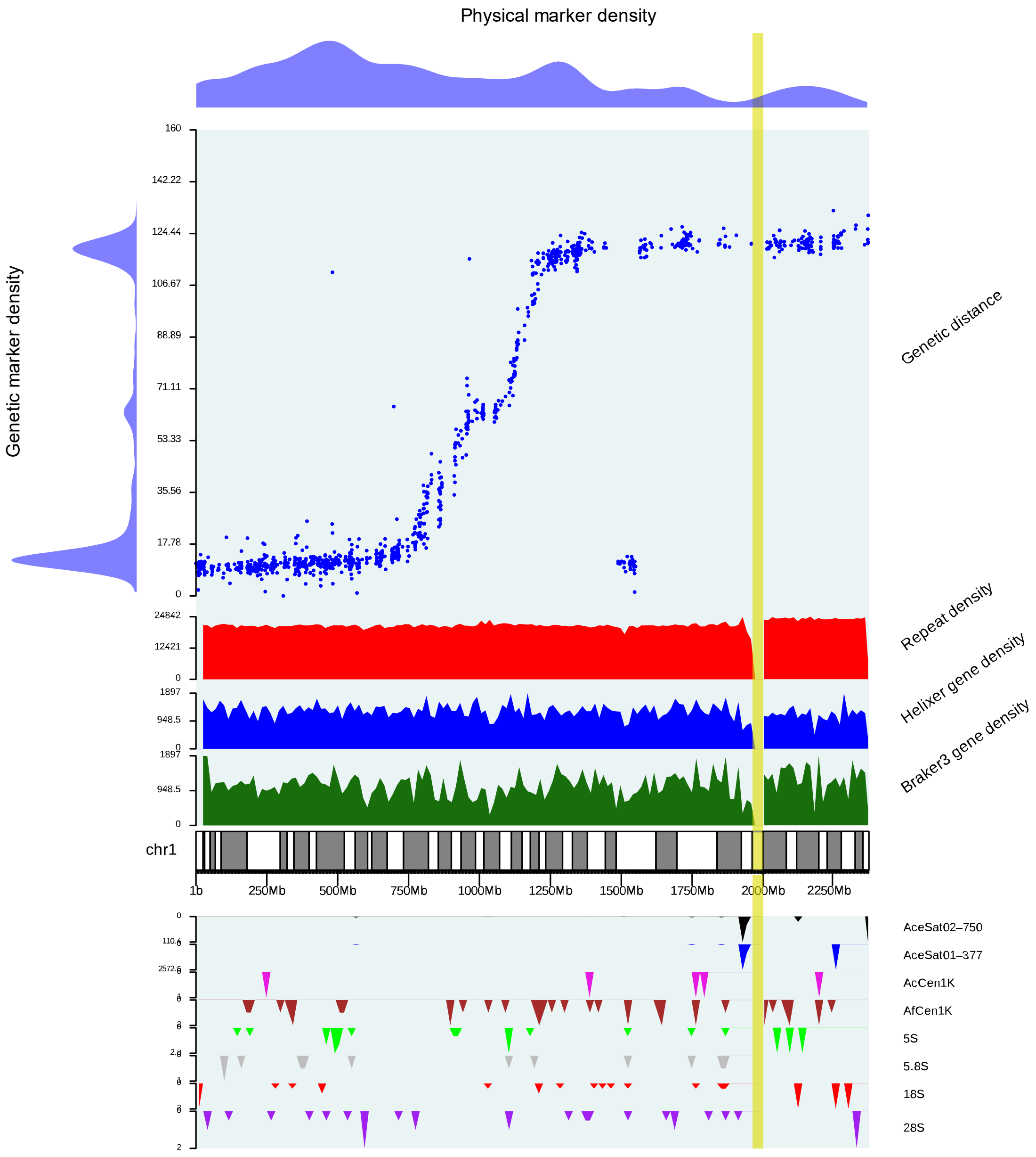

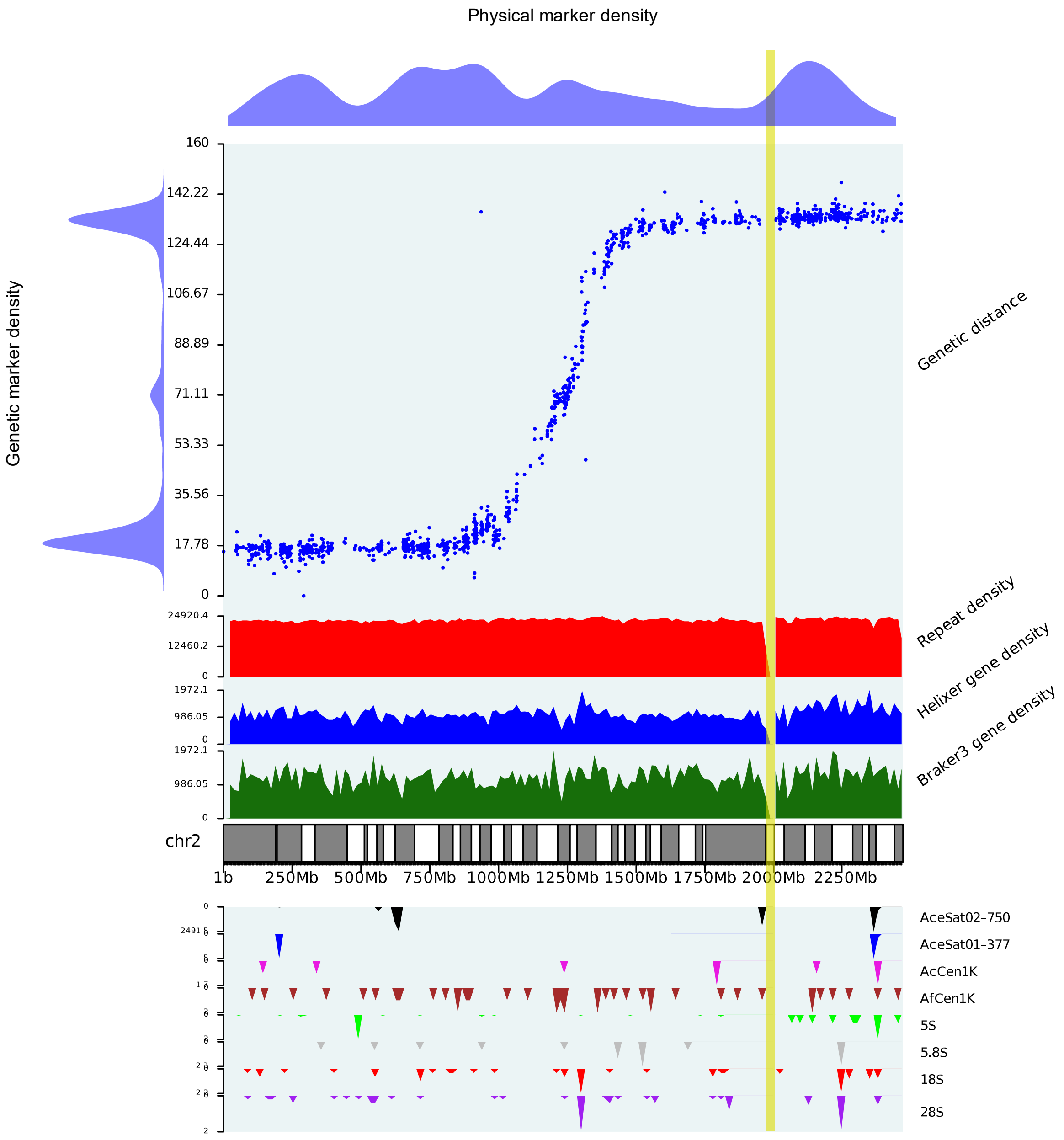

Physical marker density
Genetic marker density
Genetic distance
Repeat density
Helixer gene density
Braker3 gene density
chr2
1b
250Mb
500Mb
750Mb
1000Mb
1250Mb
1500Mb
1750Mb
2000Mb
2250Mb
AceSat02–750
AceSat01–377
AcCen1K
AfCen1K
5S
5.8S
18S
28S

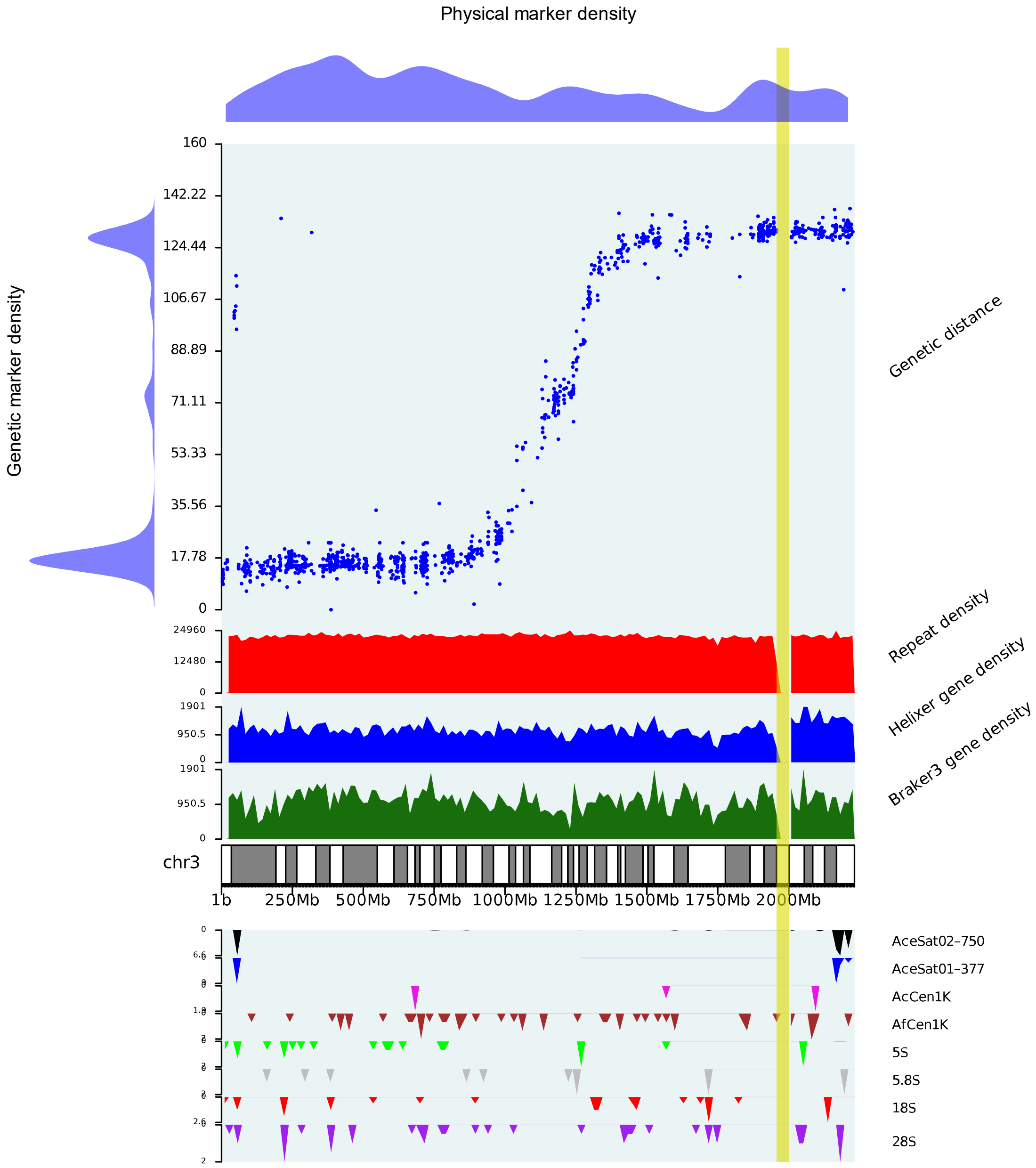

Physical marker density
Genetic marker density
160
142.22
124.44
106.67
88.89
71.11
53.33
35.56
17.78
0
Genetic distance
24960
12480
0
Repeat density
1901
950.5
0
Helixer gene density
1901
950.5
0
Braker3 gene density
chr3
1b
250Mb
500Mb
750Mb
1000Mb
1250Mb
1500Mb
1750Mb
2000Mb
AceSat02–750
AceSat01–377
AcCen1K
AfCen1K
5S
5.8S
18S
28S

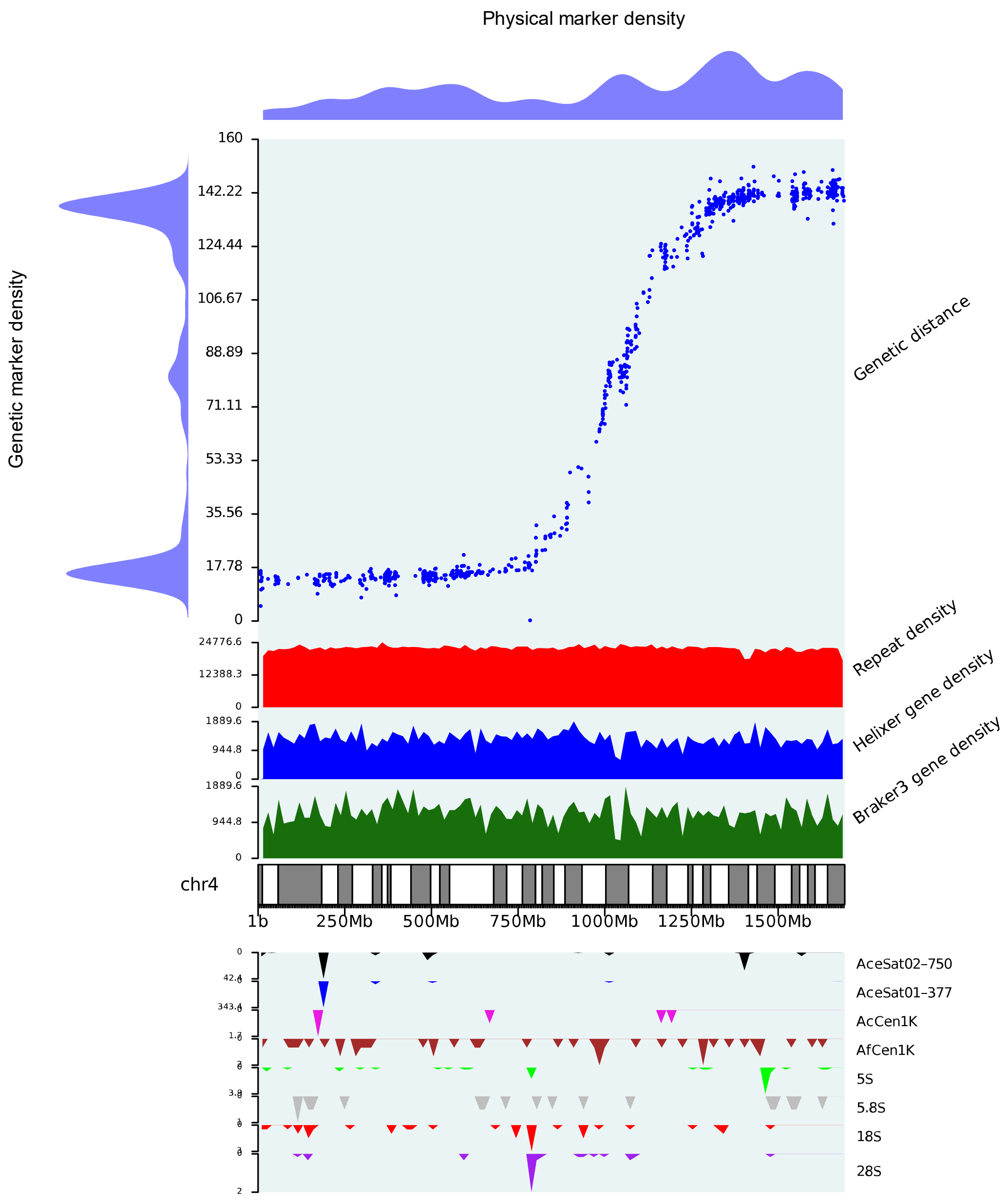

Physical marker density
Genetic marker density
160
142.22
124.44
106.67
88.89
71.11
53.33
35.56
17.78
0
Genetic distance
24776.6
12388.3
0
Repeat density
1889.6
944.8
0
Helixer gene density
1889.6
944.8
0
Braker3 gene density
chr4
1b
250Mb
500Mb
750Mb
1000Mb
1250Mb
1500Mb
AceSat02-750
AceSat01-377
AcCen1K
AfCen1K
5S
5.8S
18S
28S

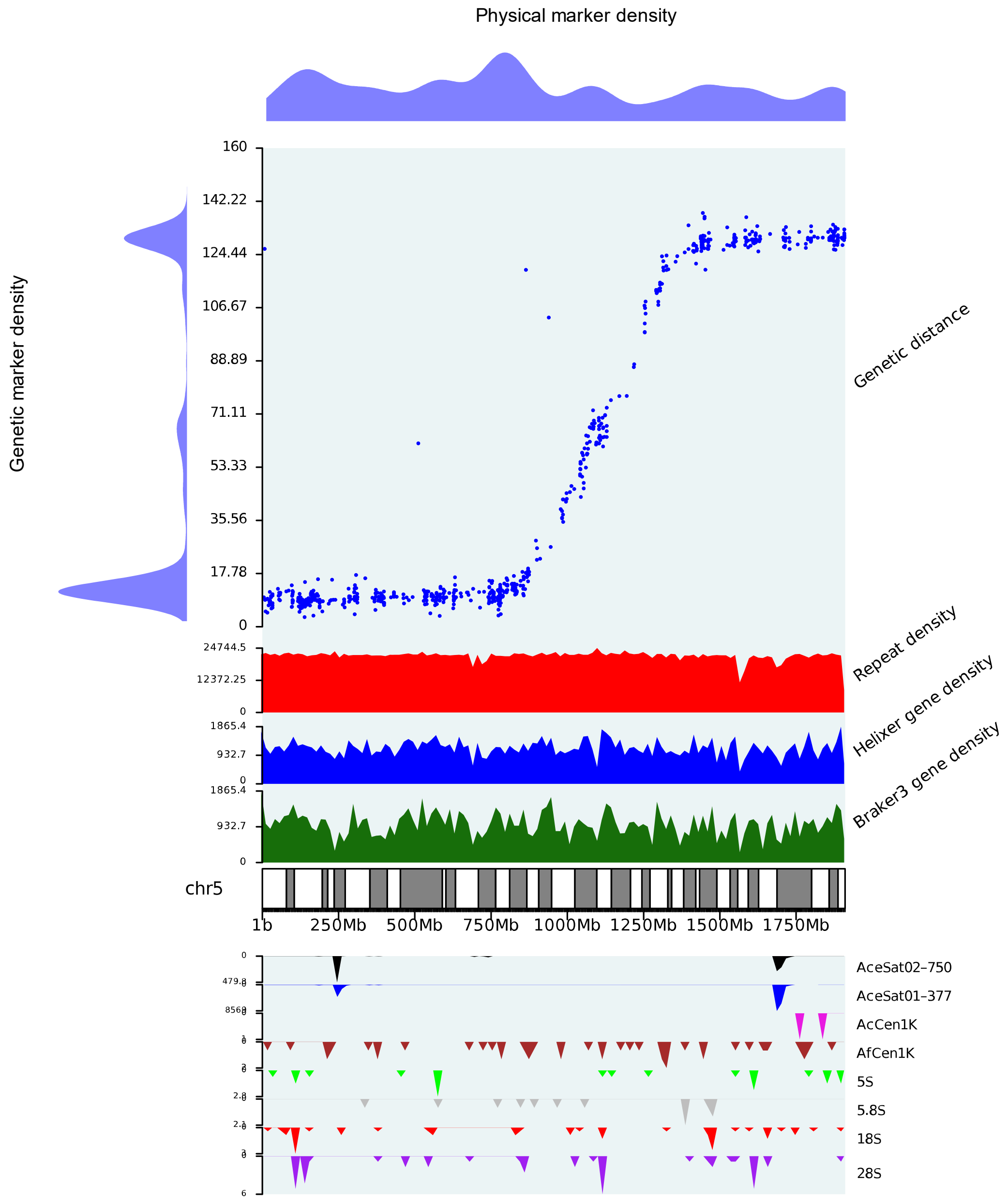

Physical marker density
Genetic marker density
160
142.22
124.44
106.67
88.89
71.11
53.33
35.56
17.78
0
Genetic distance
24744.5
12372.25
0
Repeat density
1865.4
932.7
0
Helixer gene density
1865.4
932.7
0
Braker3 gene density
chr5
1b
250Mb
500Mb
750Mb
1000Mb
1250Mb
1500Mb
1750Mb
AceSat02–750
AceSat01–377
AcCen1K
AfCen1K
5S
5.8S
18S
28S

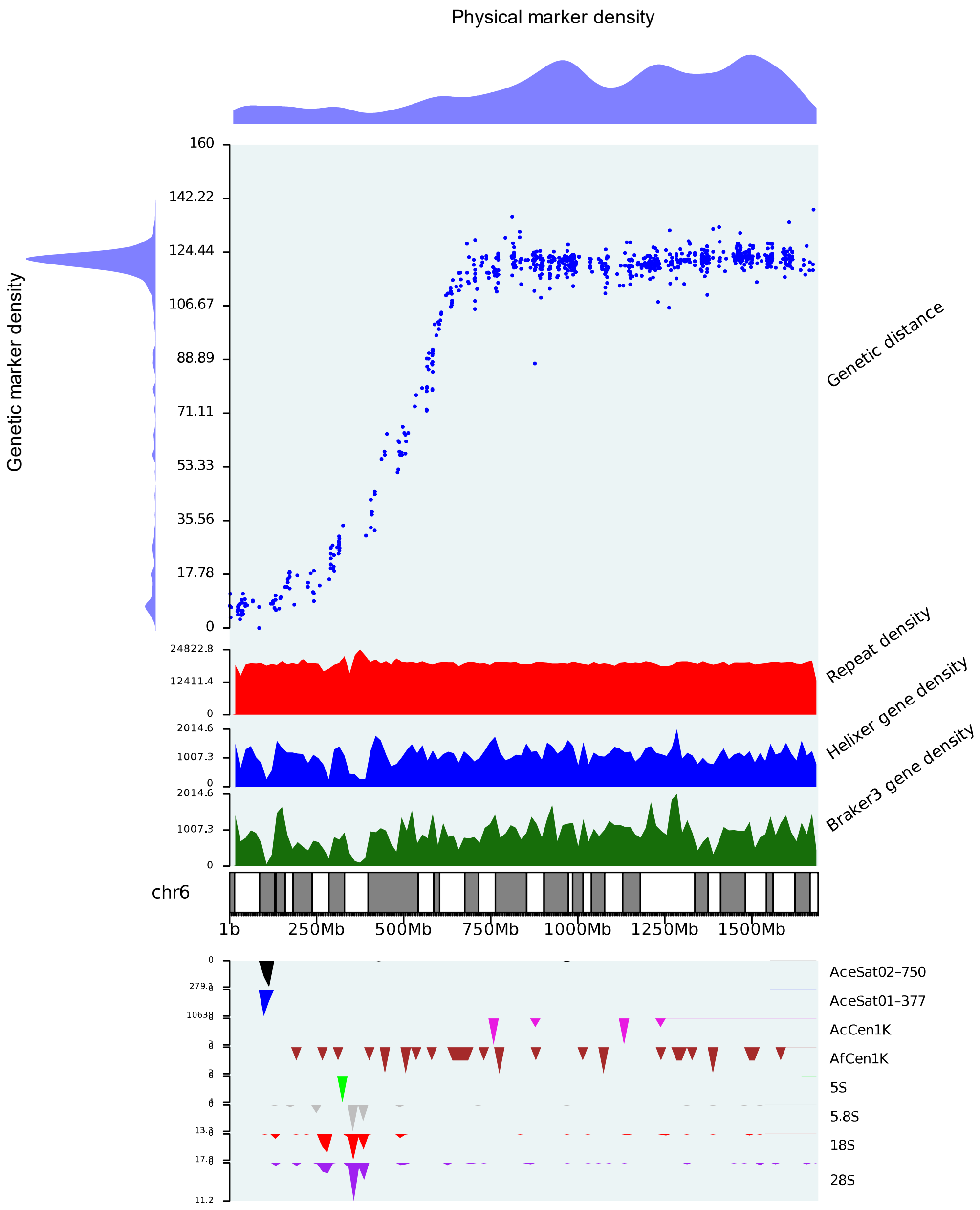

Physical marker density
Genetic marker density
160
142.22
124.44
106.67
88.89
71.11
53.33
35.56
17.78
0
Genetic distance
24822.8
12411.4
0
Repeat density
2014.6
1007.3
0
Helixer gene density
2014.6
1007.3
0
Braker3 gene density
chr6
1b
250Mb
500Mb
750Mb
1000Mb
1250Mb
1500Mb
AceSat02–750
AceSat01–377
AcCen1K
AfCen1K
5S
5.8S
18S
28S

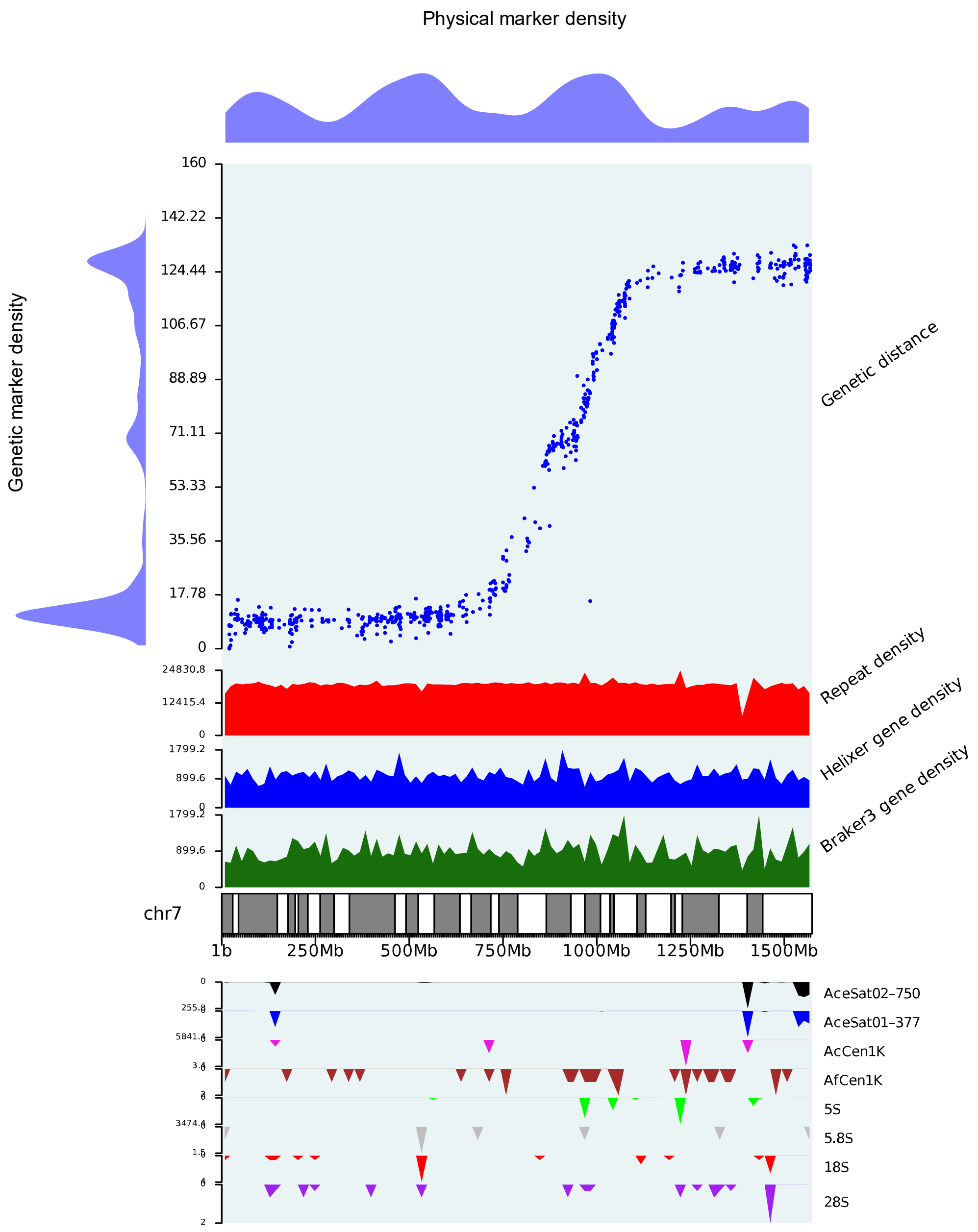

Physical marker density
Genetic marker density
Genetic distance
Repeat density
Helixer gene density
Braker3 gene density
chr7
1b
250Mb
500Mb
750Mb
1000Mb
1250Mb
1500Mb
160
142.22
124.44
106.67
88.89
71.11
53.33
35.56
17.78
0
24830.8
12415.4
1799.2
899.6
AceSat02–750
AceSat01–377
AcCen1K
AfCen1K
5S
5.8S
18S
28S

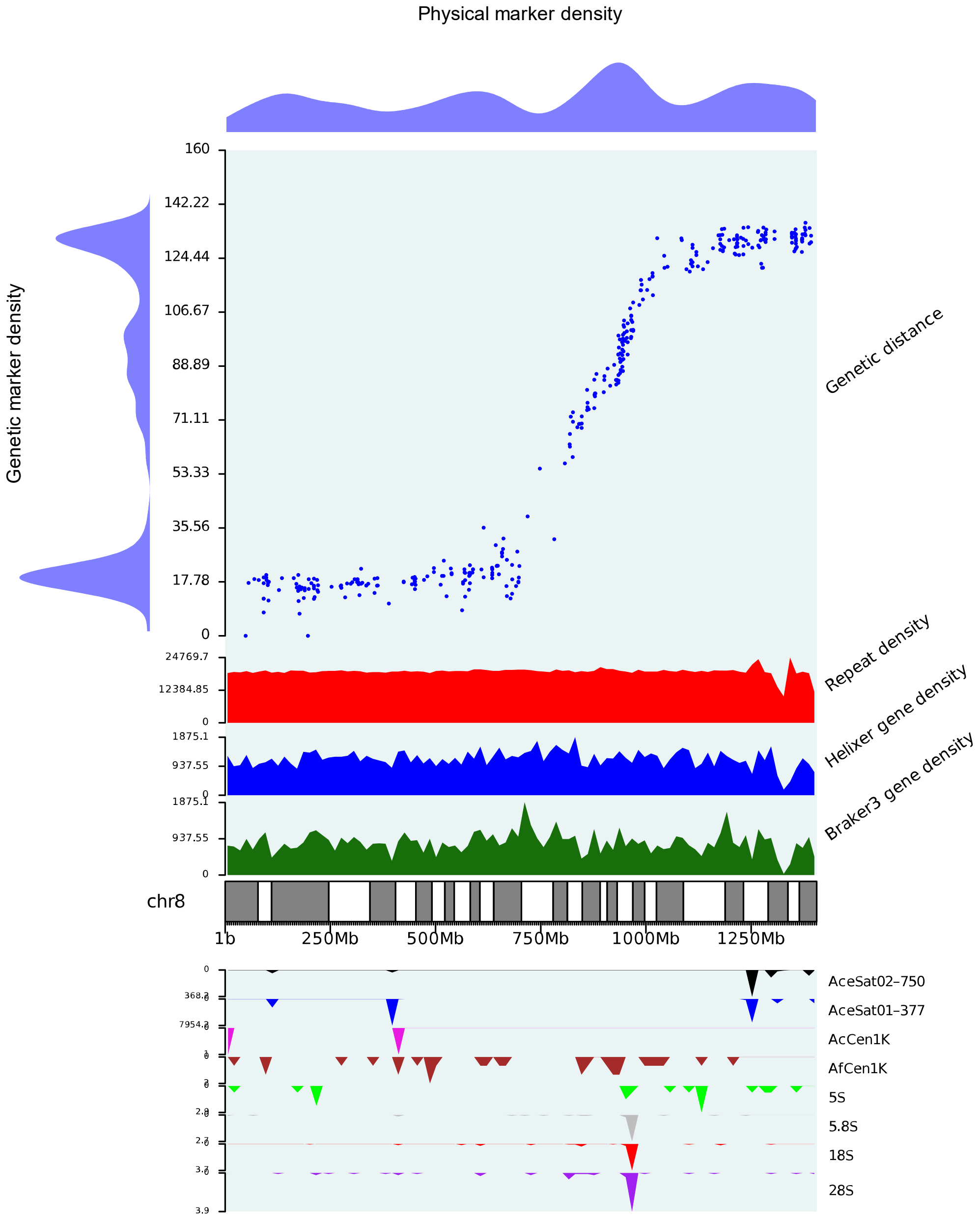
Physical marker density
Genetic marker density
Genetic distance
Repeat density
Helixer gene density
Braker3 gene density
chr8
1b
250Mb
500Mb
750Mb
1000Mb
1250Mb
AceSat02–750
AceSat01–377
AcCen1K
AfCen1K
5S
5.8S
18S
28S
160
142.22
124.44
106.67
88.89
71.11
53.33
35.56
17.78
0
24769.7
12384.85
1875.1
937.55

### S11 Physical marker mappings between A. porrum and A. sativum (2021), A. sativum (2023) & A. cepa

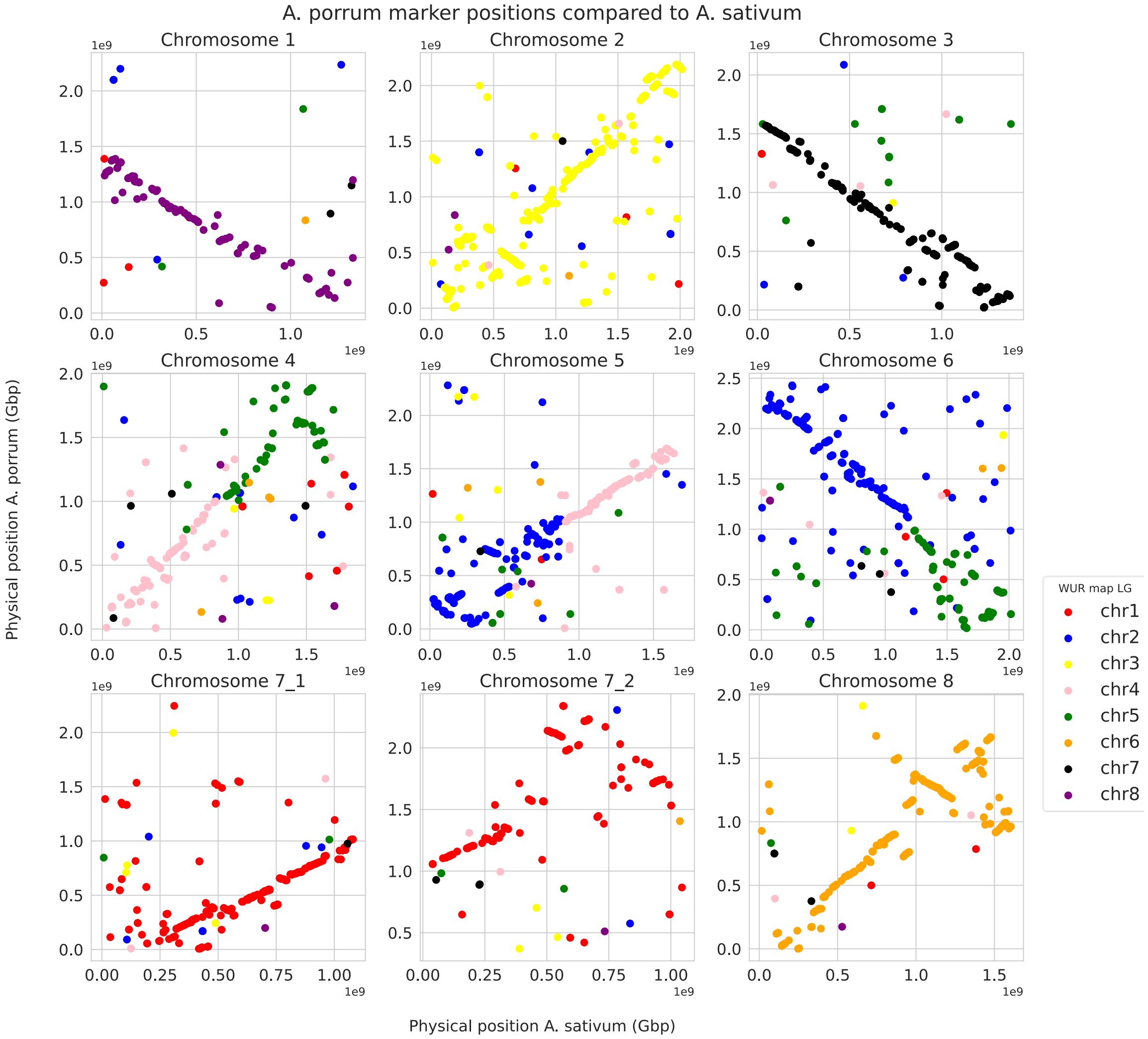

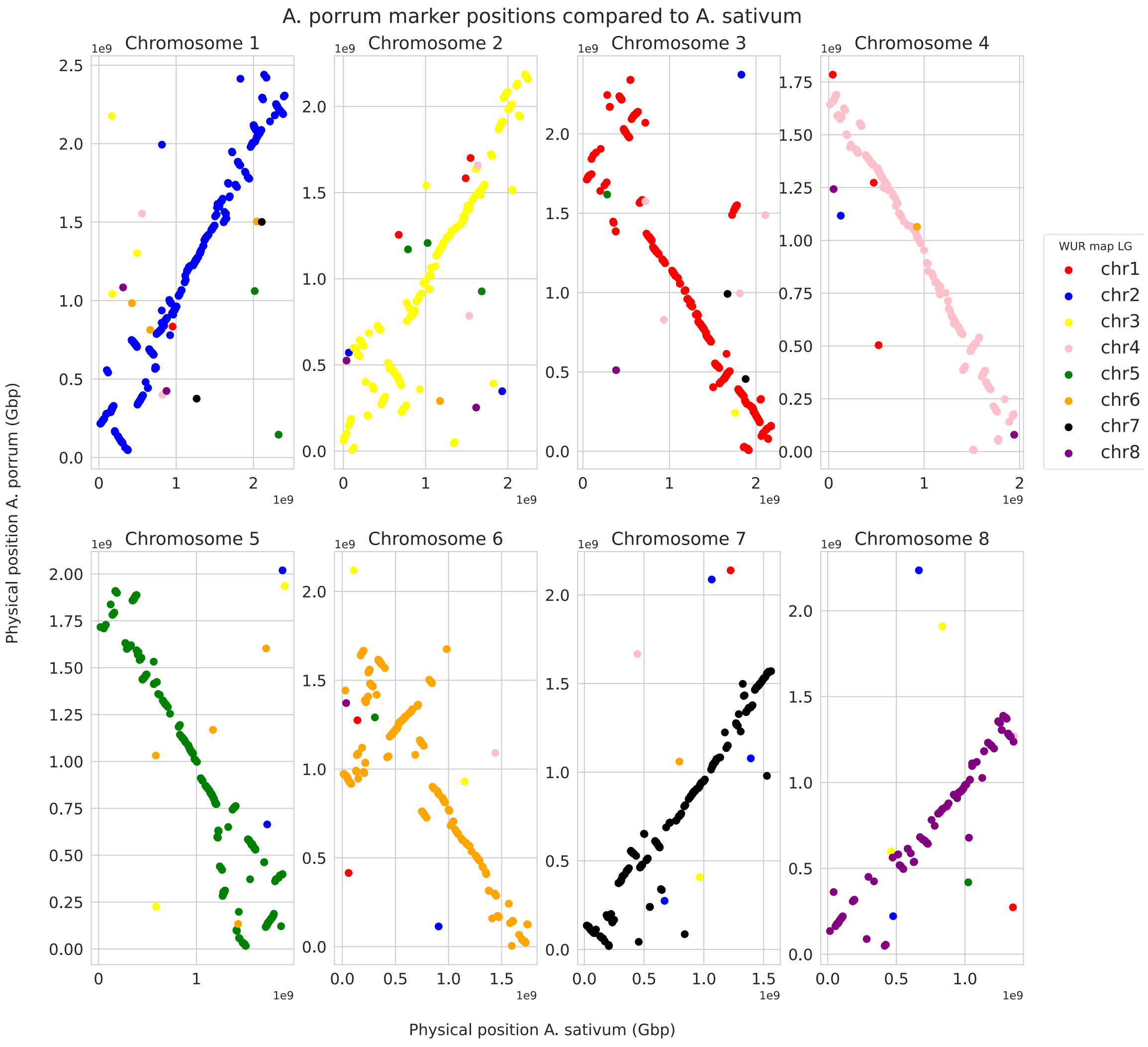
A. porrum marker positions compared to A. sativum
Chromosome 1
Chromosome 2
Chromosome 3
Chromosome 4
Chromosome 5
Chromosome 6
Chromosome 7
Chromosome 8
Physical position A. porrum (Gbp)
Physical position A. sativum (Gbp)
WUR map LG
chr1
chr2
chr3
chr4
chr5
chr6
chr7
chr8

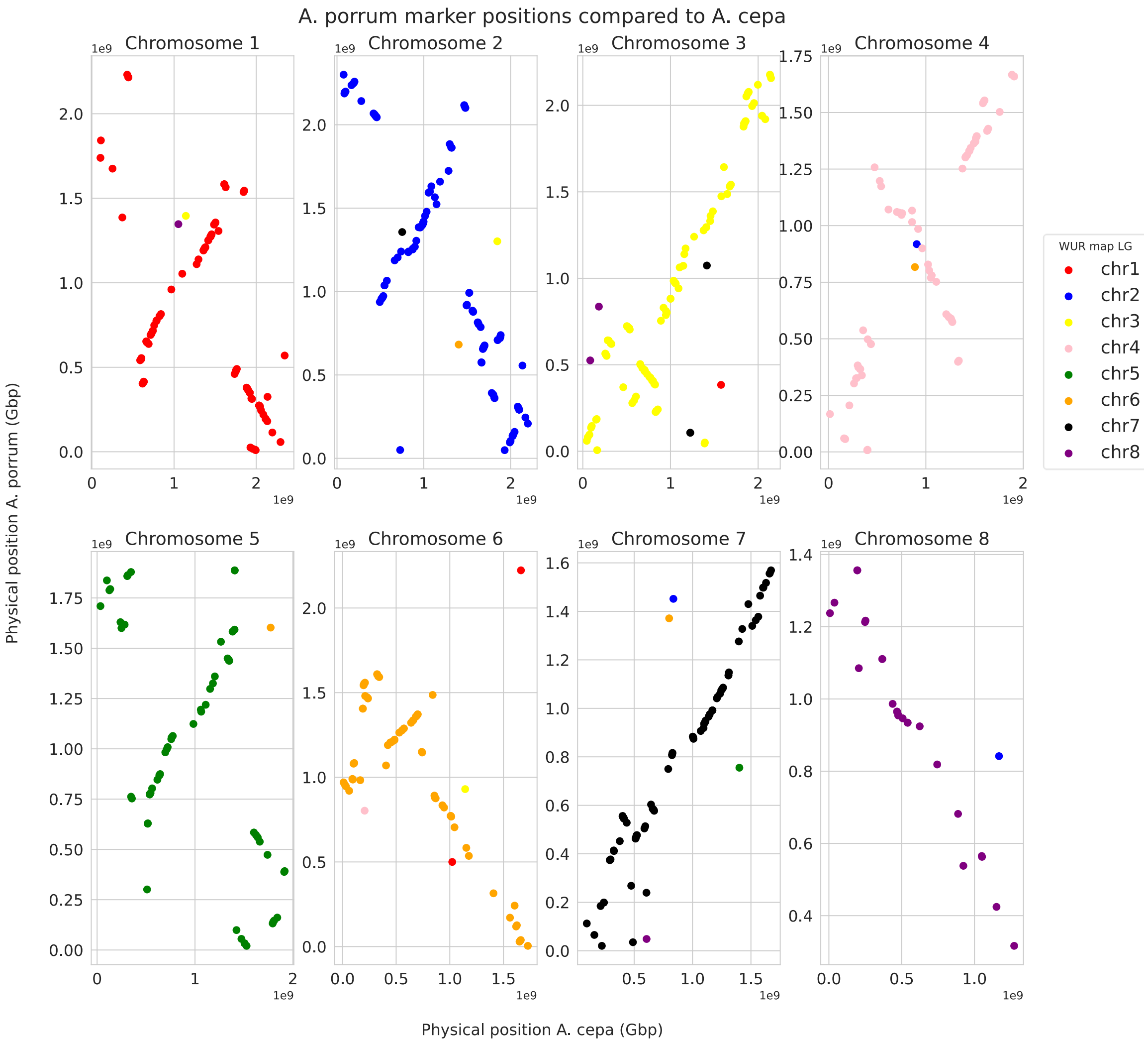
A. porrum marker positions compared to A. cepa
Chromosome 1
Chromosome 2
Chromosome 3
Chromosome 4
Chromosome 5
Chromosome 6
Chromosome 7
Chromosome 8
Physical position A. porrum (Gbp)
Physical position A. cepa (Gbp)
1e9
WUR map LG
chr1
chr2
chr3
chr4
chr5
chr6
chr7
chr8
